# Modeling Alternate Conformations with Alphafold2 via Modification of the Multiple Sequence Alignment

**DOI:** 10.1101/2021.11.29.470469

**Authors:** Richard A. Stein, Hassane S. Mchaourab

## Abstract

The unprecedented performance of Deepmind’s Alphafold2 in predicting protein structure in CASP XIV and the creation of a database of structures for multiple proteomes is reshaping structural biology. Moreover, the availability of Alphafold2’s architecture and code has stimulated a number of questions on how to harness the capabilities of this remarkable tool. A question of central importance is whether Alphafold2’s architecture is amenable to predict the intrinsic conformational heterogeneity of proteins. A general approach presented here builds on a simple manipulation of the multiple sequence alignment, via *in silico* mutagenesis, and subsequent modeling by Alphafold2. The approach is based in the concept that the multiple sequence alignment encodes for the structural heterogeneity, thus its rational manipulation will enable Alphafold2 to sample alternate conformations and potentially structural alterations due to point mutations. This modeling pipeline is benchmarked against canonical examples of protein conformational flexibility and applied to interrogate the conformational landscape of membrane proteins. This work broadens the applicability of Alphafold2 by generating multiple protein conformations to be tested biologically, biochemically, biophysically, and for use in structure-based drug design.

## INTRODUCTION

The explosion of complete sequencing for a multitude of genomes has allowed for the generation of deeper multiple sequence alignments (MSA). These MSAs are a treasure trove of information encoding co-evolution of residues that may be far apart in the linear amino acid sequence. Multiple groups have harnessed co-evolution of residues to generate distance restraints/matrices and the subsequent construction of a three-dimensional protein structure.^1^ The latest iteration, Alphafold2 (AF2), took a significant leap forward with the quality of the predicted structures it can generate.^2,3^

A database of structural models generated by AF2 for multiple proteomes has been released (www.alphafold.ebi.ac.uk). The database contains a single conformation for each protein sequence following Anfinsen’s principle that the protein’s amino acid sequence determines the native structure of the protein.^4^ However, the deposition of a single structure for each protein belies the true ensemble nature of proteins which often undergo functionally important conformational changes. The ensemble nature of most protein structures would therefore argue that a protein sequence encodes for this conformational heterogeneity. The implication is that the distance matrix derived from the MSA should contain information on this heterogeneity although at present, the general consensus is that AF2 is only able to generate a single predicted conformation.

Here, we develop a general approach to transcend the limitation that Alphafold2 generates a single conformation and consequently predict ensembles of conformations. Our work was stimulated by the modeling of the Deepmind team of multiple conformations of the multi-drug transporter LmrP (T1024) in the most recent CASP. The Alphafold team used manually curated structures in their submission.^5^ It was noted, based on the MSA and structures for LmrP homologs, that there should be more than one conformation, including an inward- and outward-open conformation. The initial runs of AF2 yielded only one conformation, inward-open, and the MSA derived distance matrix showed regions that should be close, but were in fact far apart in the computed structure.^5^ The submission to CASP XIV entailed paring the MSA and the structural templates to include only structures that were outward-open yielding the alternate conformations that became part of the CASP submission.

This study introduces a universal method for biasing the models generated by AF2. It entails replacing specific residues within the MSA (*in silico* mutagenesis) to potentially manipulate the distance matrices leading to alternate conformations. To outline, AF2 is used to generate initial models and the MSA is modified based on possible contact points in this structure, prior structural information, or regions of uncertainty within the main structure (Fig. 1). Within AF2 is an attention network that ascertains the co-evolution of amino acid residues from the MSA. Ideally, the alteration of the amino acid column to alanine or another residue turns the attention of the network to other parts of the MSA allowing for AF2 to find alternative conformations based on other co-evolved residues.

**Fig. 1:**
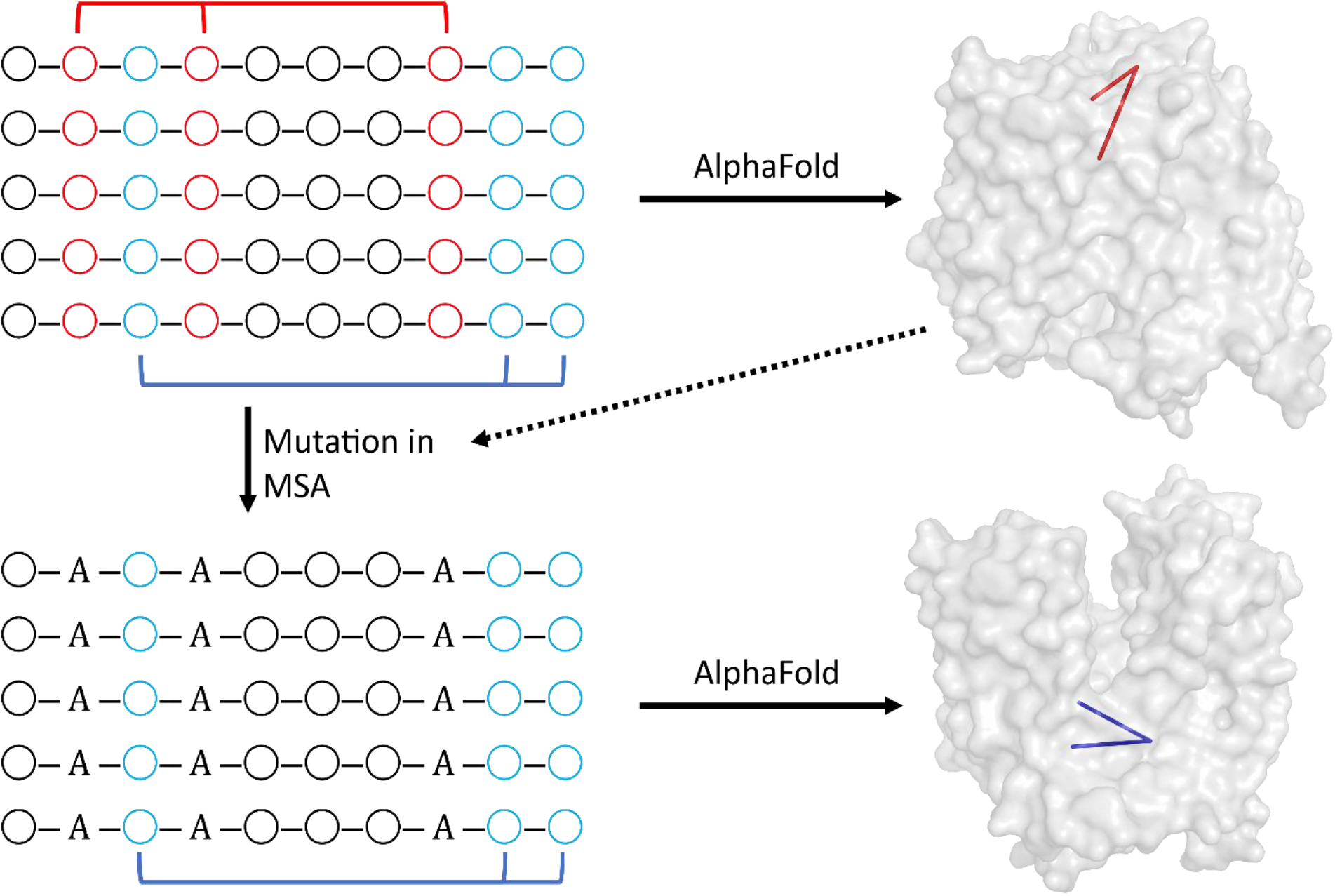
Methodology. An initially generated MSA, via MMSeqs2, is input into Alphafold2 within ColabFold to generate five structural models. Residues for mutation are chosen, in this case the three residues in red mediating a contact point on the upper surface of the protein. These mutations are made across the entire MSA (ignoring gaps). This modified MSA is then input into ColabFold for generation of new models. With the contact point in red missing, Alphafold2 within ColabFold generates a new conformation based on the contacts shown in blue.

## RESULTS

A set of protein targets were selected to illustrate the general applicability of this method and to investigate its limitations. The proteins include two classical examples of protein flexibility, adenylate kinase and ribose binding protein; LmrP a major facilitator superfamily (MFS) antiporter that was part of CASP XIV; two additional MFS proteins, XylE and GLUT5; and Mhp1 a LeuT-fold transporter. The rationale for what residues to alter in the MSA will vary depending on the protein being examined and throughout this study several methods for choosing the residues to modify in the MSA will be presented. All AF2 models/structures generated here utilize the Colab jupyter notebooks under the work termed, ColabFold.^6^

### Canonical examples of conformational flexibility

#### Adenylate Kinase

Adenylate Kinase (AK) is considered a canonical example of protein flexibility. It is a nucleoside monophosphate kinase that catalyzes the reaction ATP+AMP ⇌ 2ADP. Crystal structures of *E. coli* AK in various catalytic states have shown that this kinase undergoes large conformational changes. At the two extremes, apo AK adopts an open conformation (PDB: 4ake)^7^ whereas the inhibitor bound structure adopts a closed conformation (PDB: 1ake)^8^ (Fig. 2A).

**Fig. 2:**
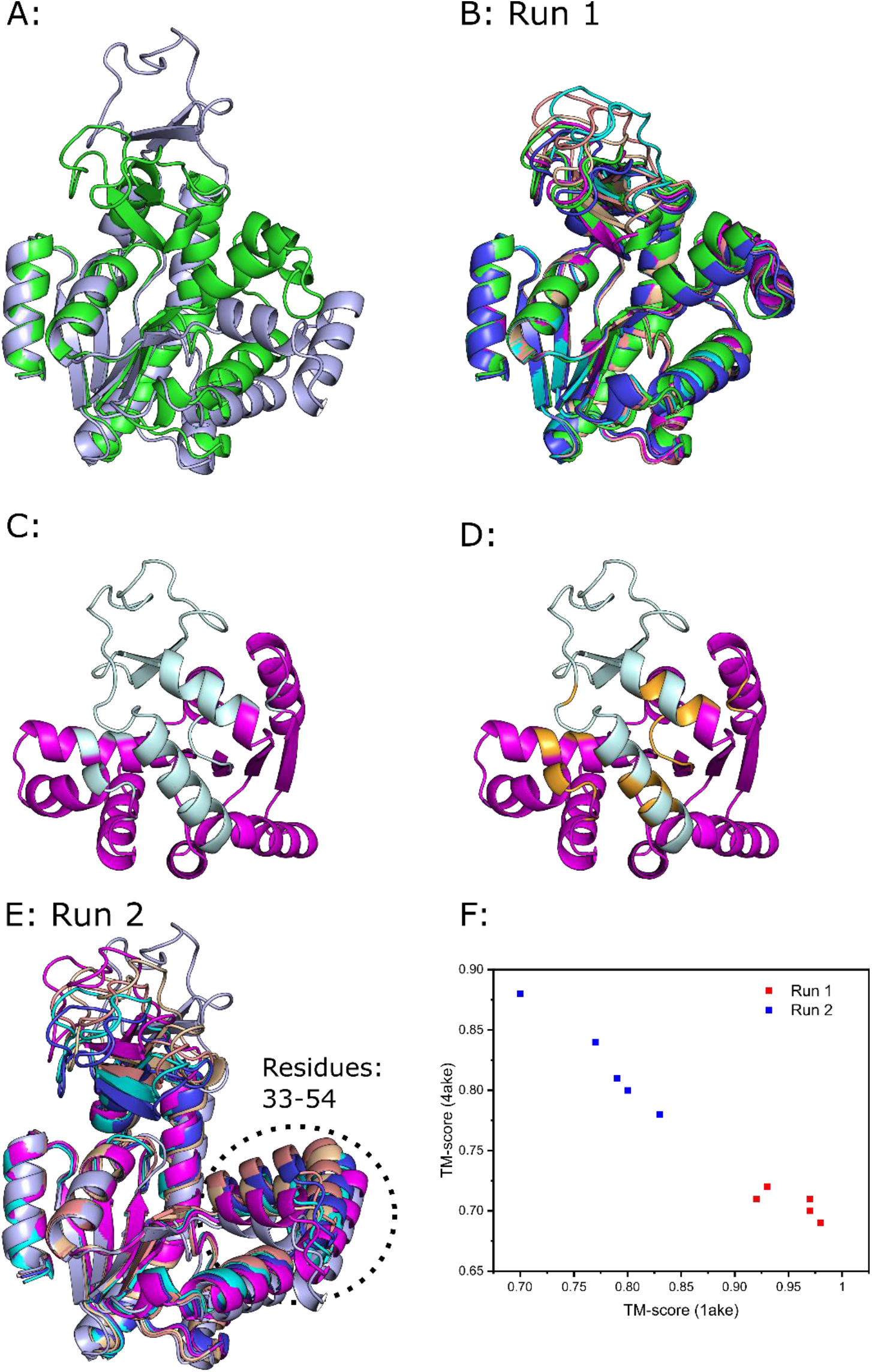
Structures of E. coli adenylate kinase. For display the structures were aligned using residues 1-25 in Pymol. A) Crystal structures for the inhibitor bound state, 1ake (green) and apo state, 4ake (lightblue). B) Five Alphafold2 models using the default MSA superimposed on 1ake (green). C) Back view of model 1 from Run 1 with the region of interest, 114-173, shown in lightcyan. D) In orange are the residues pairs that are within 4 Å of each other between the region in lightblue and the rest of the protein. E) Five Alphafold2 models after mutation of the orange residues in D (Table S1) to alanine superimposed on 4ake (lightblue). In the dashed circle are residues 33-54. F) TM-scores of the models relative to the crystal structures plotted against each other.

Input of the E. coli sequence for AK into ColabFold yields a relatively closed conformation for the five AF2 models (Fig. 2B) that are in good agreement with the x-ray crystal structure (Supp. Table 1). Based on the known ‘lid’ region of AK, the back of the kinase was chosen for potential interaction pairs and subsequent mutation to alanine (Fig. 2C-D). The MSA from the first run was altered based on these choices (Supp. Table 1) and the altered MSA and AK sequence were then used to predict new AF2 models (Fig. 2E). Although no one structure reaches the full opening of the 4ake crystal structure, the conformational flexibility is clearly manifested by the relative movement of both the ‘lid’ and the ‘flap’, residues 33-54. To compare the AF2 models to the crystal structures, TM scores were determined relative to the open and closed structures. The broad range of TM scores spanning the range between the two structures is a remarkable demonstration that this method unlocks AF2’s ability to predict alternate models describing a protein’s intrinsic conformational flexibility (Fig. 2F).

While the relative TM scores highlight the conformational flexibility, this comparison requires the availability of more than one experimental structure. Alternatively, principal component analysis (PCA) allows for a description of structural variance without multiple known protein structures. PCA for the AK crystal structures and AF2 models demonstrated that the first two components comprise 86% and 10% of the total variance of the protein. That there are two separate clusters is suggestive of more than one intermediate between the open and closed structures (Supp. Fig. 1). This small set of models is consistent with a wide variety of studies on this canonical example of protein flexibility that indicate that the ‘lid’ and ‘flap’ are capable of opening and closing independently.^9,10^

#### Ribose Binding Protein

Ribose binding protein is a bacterial periplasmic protein involved in the chemotactic response to ribose. It has also been crystallized in a number of conformations including the ones compared here: the closed structure (PDB 2dri)^11^, an open structure (PDB 1urp)^12^, and two additional open structures (1ba2 A/B) that were obtained as a consequence of the mutation, D67R^12^ (Supp. Fig. 2A).

Similar to AK, input of the amino acid sequence for the Enterobacteriase ribose binding protein yields a relatively homogeneous conformation that is most similar to the closed structure, 2dri (Supp. Fig. 2B; Supp. Table 2). As noted above, the structure with the largest opening between the two domains was with the mutation, D67R. This mutation was introduced into the MSA with the amino acid at position 67 replaced with arginine across all sequences where there is no gap. The AF2 models from this sequence and MSA yield a more diverse and open set of structures than in the absence of the mutation (Supp. Fig. 2C). This suggests that this method of modifying the MSA and input sequence is capable of directly examining structural consequences of single point mutations.

To further examine the conformational flexibility of ribose binding protein, residues mediating various connections between the two domains were mutated to alanine alone and in conjunction with the D67R mutation (Supp. Table 2). Each run yielded a diverse set of structures (Supp. Fig. 2D-I). None of the runs produced a conformation as open as the most open structure, 1ba2A (Supp. Table 2). This is not surprising as this structure was presumed to arise in large part from crystal packing.^12^ That AF2 could not generate the most open structure suggests that it did not just learn crystal structures in the database, but does generate structural models based on physical principles.

Carrying out PCA on the 8 runs of AF2 and 4 crystal structures of ribose binding protein yields percentages of 88% and 8% for the first two components. The plot of the two components against each other suggests that there might be an outlier in the structures, Run 8-5 (Supp. Fig. 3A). This is supported by the TM and RMSD values for this model relative to all four crystal structures (Supp. Table 2). PCA analysis excluding this model yields a shift in relative amount of variance for the two components, 94% vs 2%, though the resultant plot of the two values against each other is similar to the one with all the structures (Supp. Fig. 3B). Interestingly, the structure 1ba2A is outside the grouping of the rest of the structures, in line with it being considered a crystal contact artefact.^12^

### Predicting the conformational ensembles of membrane proteins

#### LmrP

LmrP, a proton-drug antiporter from *Lactoccoccus lactis*, is a member of the major facilitator superfamily (MFS) that generally couples ion gradients to transport a variety of substrates. The MFS fold typically consists of 12 transmembrane helices with two-fold symmetry between helices 1-6 and 7-12. Based on other MFS structures, spectroscopic data, and proposed catalytic cycles, LmrP should sample both inward- and outward-facing conformations.^13,14^ The crystal structure for LmrP was obtained in the presence of substrate and is outward-open.^15^

Input of the LmrP protein sequence into ColabFold yields inward facing conformations with small variations at the intracellular vestibule (Fig. 3A) in agreement with the initial models from Deepmind.^5^ To obtain an alternate outward facing conformation, the Deepmind team parsed the MSA and limited the structural templates to those that are outward facing. In contrast, the approach here does not use structural templates. Rather, the residues that mediate the interface between the two halves of the protein in the inward facing AF2 model were mutated to alanine (Supp. Table 3). These mutations prompted AF2 to output an outward conformation with small variations at the extracellular vestibule in agreement with the crystal structure (Fig. 2B).

**Fig. 3:**
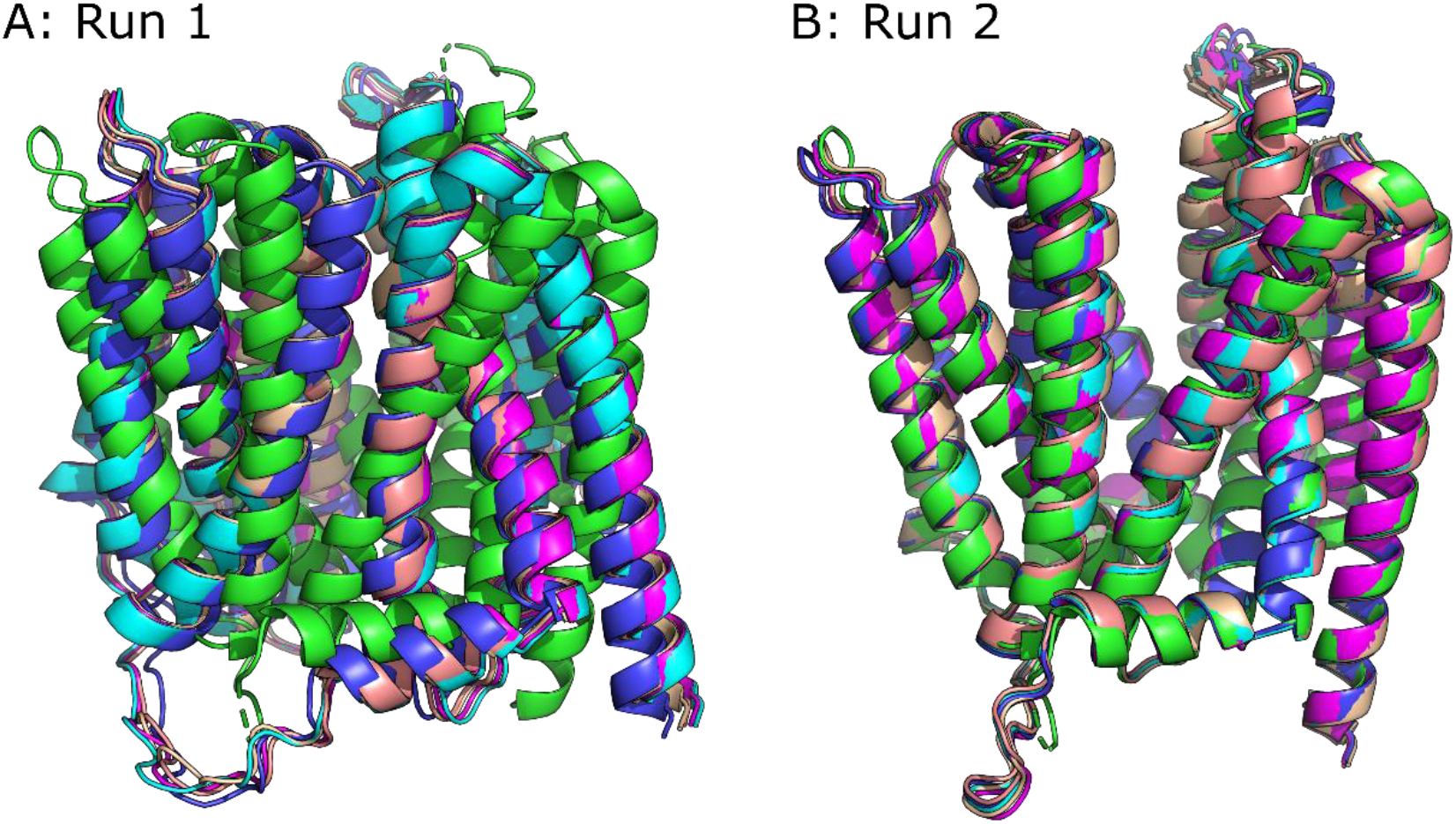
Structures of LmrP. A) Five Alphafold2 models with the default MSA superimposed on the LmrP crystal structure, 6t1z (green). B) Five Alphafold2 models after mutation of residues at the interface of the inward facing structure from the top model in A superimposed on the LmrP crystal structre, 6t1z (green).

While the two-fold symmetry of LmrP directed the initial choice of residues to alter at the interface, additional sets of mutations were also pursued. Mutation of only one half of the interface residues, either the N-terminal or C-terminal half (Supp. Table 3) lead to outward open conformations with variable positioning of the extracellular parts of the helices (Supp. Fig. 4A-B). One of the most interesting sets of mutations involved 3 residues at the center of the transmembrane region (Supp. Fig. 4C, Supp. Table 3). The resulting set of AF2 models consisted of two inward-open and three outward-open conformations, as if the models are “isoenergetic” and mimicking the interconversion of the antiporter in the presence of ligands as expected.^13,14^

To illustrate the relationship between the crystal structure and the twenty-five AF2 models, we analyzed the extent of opening by measuring two distances, one on the intracellular and one on the extracellular side of the transporter (Supp. Fig. 4E). The intracellular side (red circles) is either closed or over 70% open; while the extracellular side is slightly more diverse, with several models more open than the crystal structure (Runs 4-5, 5-2, and 5-3) and one model that is partially occluded (Run 5-4).

These structures were then analyzed by PCA, yielding one main component comprising 95% of the variance in the structures. Plotting the first two components against each other (Supp. Fig. 5) suggests that the protein rocks between the open and closed structures through a common movement of the two domains, consistent with the canonical model of MFS alternating access.

#### XylE

XylE, also a member of the MFS, is an *E. coli* xylose:proton symporter and is a close bacterial homolog of eukaryotic glucose transporters. It has been crystallized in three conformations (Supp. Fig. 6A):, inward open (PDB: 4ja4)^16^, inward occluded (PDB: 4ja3)^16^, and outward open (PDB: 6n3i)^17^, the latter was obtained by introducing a double mutation, G58W/L315W.

Input of the protein sequence into ColabFold yields a single, inward open conformation that is most similar to the inward open crystal structure (Supp. Fig. 6B). Mutating the residues that mediate the interaction between the two halves of this MFS leads to a switch to the outward-open conformation with some variability in the ends of the helices on the extracellular side (Supp. Fig. 6C, Supp. Table 3). Similar to LmrP, a slightly more open conformation was observed when the C-terminal domain residues of the interface were mutated, though for model 3 the two halves of the protein are no longer in contact (Supp. Fig. 6D, Supp. Table 3). This distorted structure is most likely a result of removing too many key contact points between the two halves of the protein. The presence of this distorted structure suggests that the other structures from this run should be interpreted with caution in any subsequent analysis as the reduction in contact points may be too large and over biasing the other models as well. Using only the N-terminal domain residues of the interface leads to outward-open conformations with slightly more variability in the ends of the extracellular helices (Supp. Fig. 6E).

As noted above, the outward-open crystal structure, 6n3i, was obtained with two point mutations. Mutation of these two residues across the MSA also leads to an outward open conformation (Supp. Fig. 6F) and in fact yields the best matching conformation to 6n3i by TM score (Supp. Table 4). The bias towards an outward open conformation along with the results above suggest that AF2 can model the effects of specific mutations, though the broad applicability of this observation needs to be rigorously established.

Shown in Supplemental Figure 6G is the top model from the initial, unbiased AF2 prediction (Run 1) colored by pLDDT score. The pLDDT score is Alphafold’s metric for ranking the confidence in the structure at every residue. Of note is the less confident regions on the intracellular ends of the helices (lighter color blue). This could reflect uncertainty of the exact position of these regions, potentially indicative of conformational heterogeneity. With that in mind, two less confident regions (154-173 and 327-341) were chosen as regions for potential interactions. Mutation of the selected residues and their partners within 4 Å did not switch the transporter to an outward facing state, though there is some alteration of the conformation compared to the structures in the unbiased run (Supp. Fig. 6H-I). In fact, each of the transporters were less open on the intracellular side than without the mutations (Supp. Fig. 6J), indicative of an occluded conformation.

The structures generated here for XylE yield a more complex set of principal components than LmrP with the first three components having percentages of 56%, 15%, and 10%. Plots of the first component vs the second and third components suggest that there might be outliers leading to the high variability of the structures (Supp. Fig. 7A). One potential outlier, Run 3-3, is the misfolded model where the N- and C-terminal halves were dissociated. While the plot of the first vs the third component suggest that all of Run 2 might be outliers only Run 3-3 and Run 2-1 were removed for the next analysis. This led to changes in the populations of the components to 71%, 10%, and 6%. Comparing principal components 1 vs 2 confirms that the rest of Run 2 could be outliers relative to the rest of the structures (Supp. Fig. 7B). Removing these from the analysis shifts the percentages of the first three components to 82%, 6%, and 2%. The plot of the first two components has a similar shape as Lmrp (Supp. Fig. 7C) suggesting that the protein should oscillate through a common movement of the two domains.

#### GLUT5

While belonging to the MFS, GLUT5 is a mammalian fructose facilitator, unlike LmrP and XylE. As a facilitator it only transports fructose down its concentration gradient. It has been crystallized in two conformations, outward-open (PDB: 4ybq) and inward-open (PDB: 4yb9).^18^

Input of the protein sequence into ColabFold leads to two conformations (Fig. 4A). This differs from the other proteins shown so far that yielded only a single conformation with some small variability between them. The potential rationale for this diversity could be that this protein acts in facilitative diffusion so there is less preference for a specific conformation and this is reflected in the MSA and Alphafold’s decoding of the co-evolved residues. Although no manipulation of the MSA was needed to generate the two conformations, the web database for the human proteome predictions returns only one conformation, supporting having more than one structure in the databank when the five AF2 models do not have the same conformation.

**Fig. 4:**
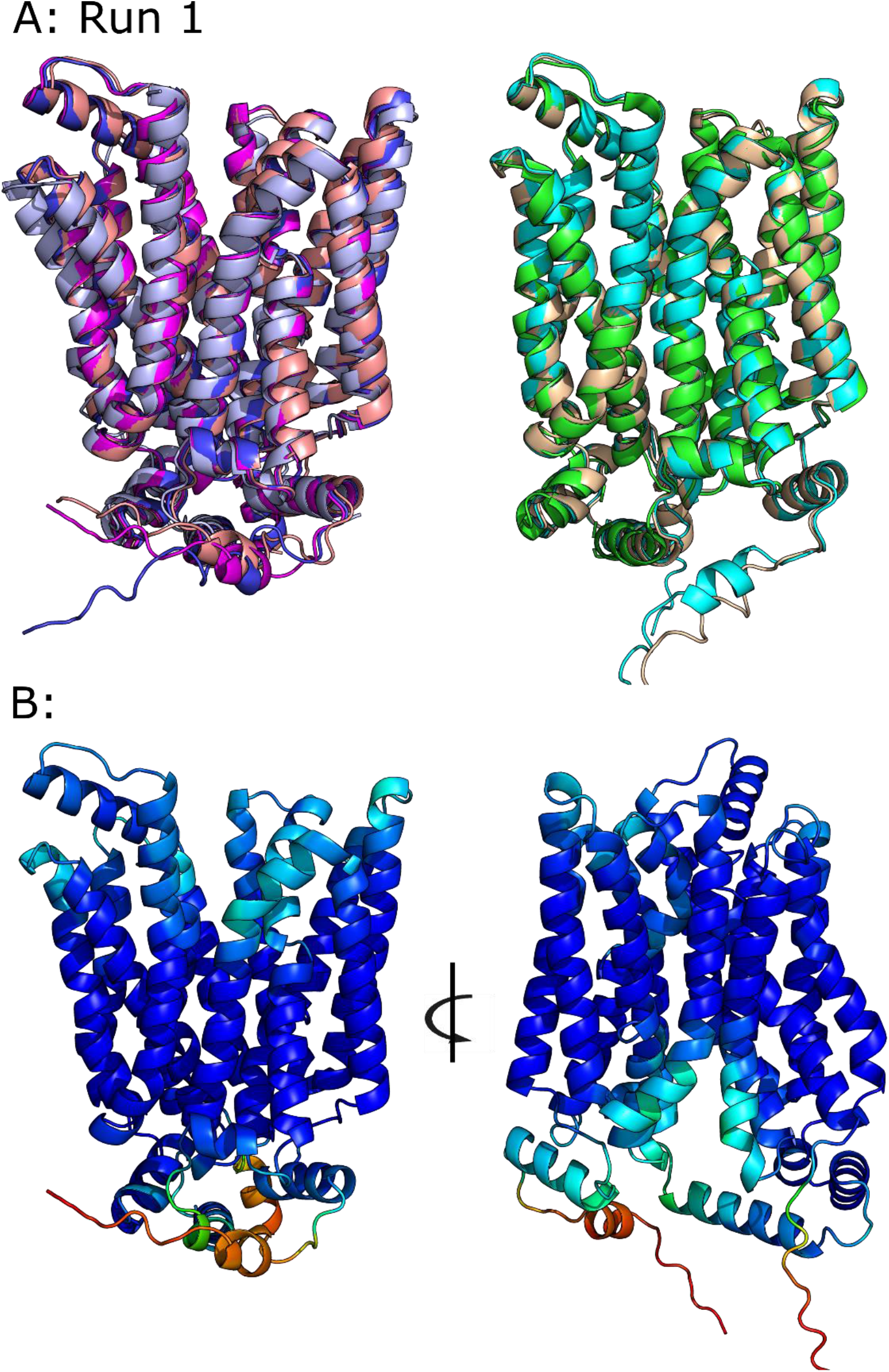
Structures of GLUT5. A) Models from the default Alphafold2 run. Left: three Alphafold2 models superimposed on the outward open structure, 4ybq (light blue). Right: two Alphafold2 models superimposed on the inward open structure, 4yb9 (green). B) Top models for each conformation colored by pLDDT score highlighting the lower score on the open side of the proteins. Left: the outward open model rotated 180°. Right: the inward open model.

Plots of the pLDDT values for the top model for each conformation indicate uncertainty in the structure for each orientation (Fig. 4B). The location of the uncertainty is striking as they are at the ends of the helices on the open sides of the protein, similar to LmrP (Supp. Fig. 4D) and XylE (Supp. Fig. 6G). While these regions of uncertainty are suggestive of conformational heterogeneity, it does not imply that these are the direct regions of flexibility. It is likely that these regions have elements in the distance matrix that are not being satisfied in the generated protein structure and there is an alternate conformation that would satisfy these unused elements of the distance matrix.

Similar to the other MSF proteins examined, the interacting residues in the top model for the inward and outward facing structures were determined (Supp. Table 5). The number of residues that are interacting in the GLUT5 models are larger than in the others. This could be due to the tighter interface of the facilitator compared to active transporters. Nevertheless, running the modified MSA’s through AF2 yielded the anticipated alternate conformations (Supp. Fig. 8A-B).

Principal component analysis of GLUT5 yields the first three components having 66%, 11%, 8% of the variance in the structures. Plots of the first vs the second or third component are not similar to LmrP or XylE (Supp. Fig. 9). The outward open structure, 4ybq, based on the second component would appear to be an outlier as there are no other structures with this movement. Alternatively, this particular conformation may not have been explored by the generated AF2 models. Whether 4ybq is an outlier or GLUT5 has a unique conformational distribution relative to the other MFS transporters is uncertain, though the difference could be a consequence of mechanistic divergence between the facilitator GLUT5 and the secondary active transporters LmrP and XylE. The difference in the principal components compared to XylE, if fully supported by additional analysis, would suggest an alternate conformational cycle for GLUT5.

#### Mhp1

The final target Mhp1, a benzylhydantoin transporter from *Microbacterium liquefaciens*, was selected not only because it belongs to one of the largest fold class of transporters, the LeuT-fold, but also because of its apparent structural heterogeneity. It has been crystallized in three conformations: outward open (PDB: 2jln)^19^, substrate bound, closed (PDB: 4d1b)^20^, and inward open (PDB: 2×79)^21^ (Supp. Fig. 10A) without the need for mutation or conformationally-selective antibodies. Isomerization between the three structures involves shifts in helices 9/10, the extracellular loop 2 (EL2), and the N-terminal portion of helix 5. Spectroscopic analysis confirmed the ligand-dependent transition of Mhp1 between the three conformations.^22^

Input of the Mhp1 sequence into Colabfold yields more than one conformation with all three crystal structures being represented by one of the models (Supp. Fig. 10B). Examination of the pLDDT values indicates that the regions with the largest uncertainty correspond to the regions of diversity between the crystal structures (Supp. Fig. 10C). This supports the notion that these regions of uncertainty potentially have unsatisfied distances in the distance matrix derived from the MSA.

Residues 159-190 of TM5 was the first region targeted for alanine mutagenesis (Supp. Table 6). The five AF2 models are highly consistent and are similar to the closed structure (Supp. Fig. 10D; Supp. Table 6). Instead of mutating the residues that span between 159-190 and the rest of the protein, only the residues that are within 4 Å of 159-190 were mutated. This would potentially allow for the residues in TM5 to find new contact sites to the rest of the protein. Again, the five AF2 models are highly consistent, but now the structures take on elements of the inward-open structure (Supp. Fig. 10E; Supp. Table 6). The next region examined is the variable region that spans TM 9/10 and the subsequent AF2 models have slightly more variability compared to the previous runs, and the structures now take on elements of the closed- and outward-open structures (Supp. Fig. 10F; Supp. Table 6). While modification of the MSA does yield additional conformations, the conformations most like the crystal structures were obtained on the initial, unmodified run. This further supports including multiple structures in the AF2 structural database and highlights the need to obtain all five AF2 models as the modelers are not monolithic.

PCA of Mhp1 indicates that the movements of the protein occur along multiple axes as the first three components comprise 56%, 26%, and 4% of the conformational variance. The components are consistent with the complex movements seen in the crystal structure. The plot of the first two components indicates that the three crystal structures and the initial AF2 models appear to segregate along one line, while the additional models appear to segregate along another line (Fig. 5). The simplest transport cycle for the symporter Mhp1 should transit through 4 states, outward-open → bound-occluded → inward-open → empty-occluded → outward-open. With the biochemical context of the crystal structures, tracing a potential path through the three structures would suggest that the lower path with the new alternate conformations captures the protein returning to the outward-open state through previously unseen empty-occluded conformations.

**Fig. 5:**
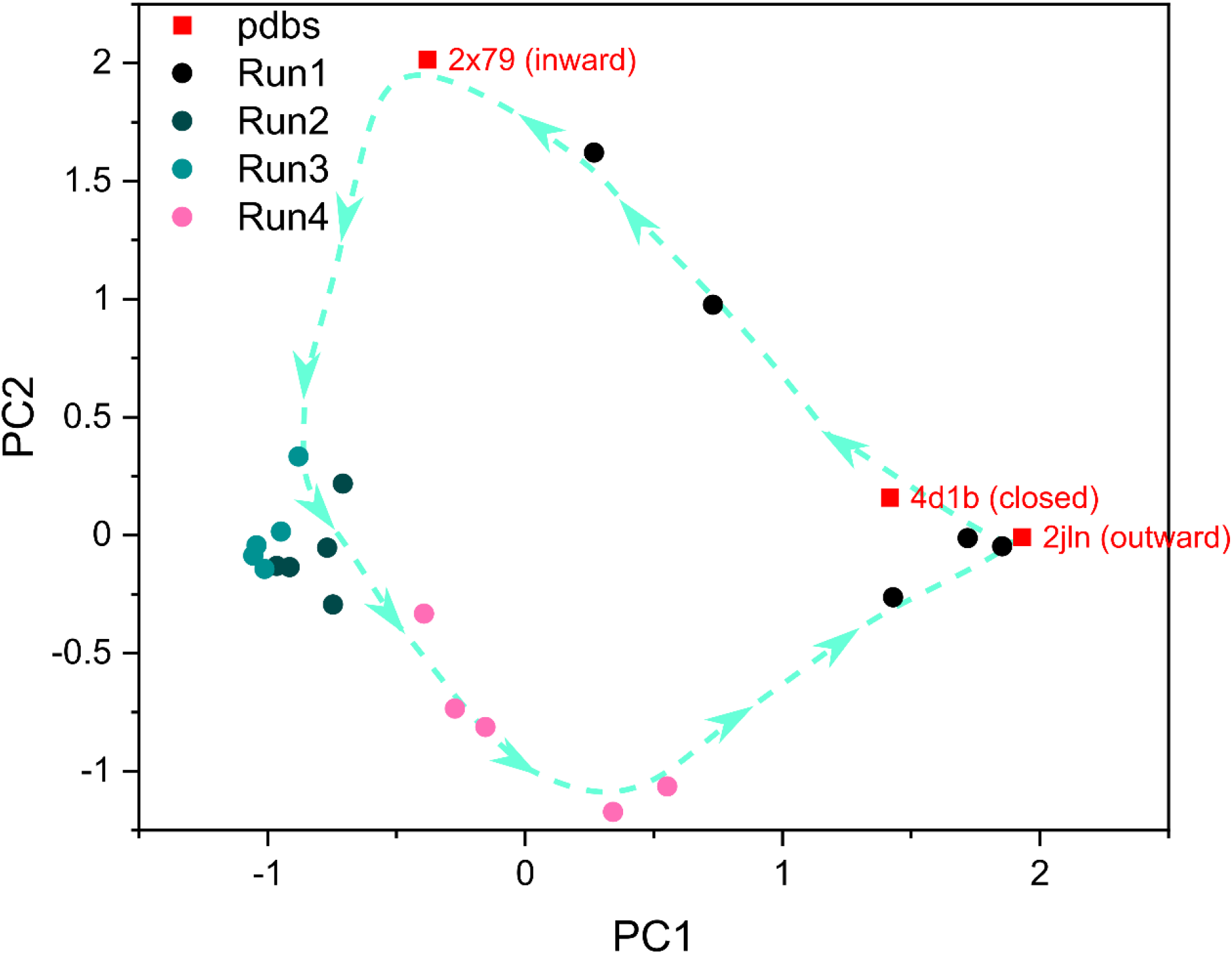
Plot of the first two principal components for the structures from the four runs for Mhp1. In red are the three crystal structures with each run colored independently given in the figure. The dashed line is a putative reaction cycle with arrows denoting the path. Starting at 2jln (outward) leads through the closed structure going through 2 of the Run 1 structures toward the 2×79 (inward) structure. The “new” models for the runs exploring contacts in TM5 and TM9/10 would comprise the return of the protein through an alternate pathway of a second closed structure back to the outward facing structure.

## DISCUSSION

The premise of the work presented here is that the MSA contains information on multiple protein conformations. Therefore, it follows that AF2 can generate these conformations with appropriate modifications of the MSA. Our method entails simple *in silico* mutagenesis to successfully coerce AF2 to sample alternative conformations for a number of target proteins both water-soluble and membrane embedded (Figs. 2-4, Supp. Figs. 2, 4, 6, 8, 10). For Ribose Binding Protein, AF2 was not able to generate models consistent with one of the crystal structures supporting the idea that AF2 does not just remember structures, but is able to generate multiple conformations rooted in physical principles.

In a recent preprint, a different methodology for sampling protein conformational space with AF2 was described.^23^ To obtain multiple conformations several alterations in the AF2 pipeline were made, there was no recycling within the AF2 module and the number of sequences of the MSA seen by AF2 was modified. In addition, the set of 8 protein targets had neither of the structures describing the conformational change in AF2’s training set. The authors were able to obtain both conformations in 7 targets without templates and in the last case with templates. They also examined 4 cases where one of the structures was in AF2’s training set, but for these the methodology was unable to generate an alternate conformation.

To compare these two methods, four of the proteins from that study are examined here. Two of the proteins had no structures in the training set, LAT1 and MCT1, while two proteins MurJ and PfMATE only had one of the two conformations in the training set. The changes to the MSA are given in the Supplemental Information. Plots of the TM-score for the AF2 models generated by making *in silico* mutations in the MSA suggest that the approach described here is capable of generating both conformations for all targets including those where the alternative method failed (Supp. Fig. 11). This demonstrates the general utility of our methodology in generating ensembles of multiple conformations regardless of whether the protein is in the training set or not.

The results presented here suggest that in some cases single mutations can lead to structural changes in the models generated by AF2 (Supp. Figs. 1 and 3). Recent reports have utilized AF2 models and existing paradigms to ascertain the potential effect of a mutation without a discernible correlation.^24,25^ Here the structures of the mutant protein lead to an increase in disorder or altered conformations. The challenge will be in devising metrics to quantitate and evaluate the effect of the mutation on the structure relative to the wild type structure. The metrics would then be compared to functional data to test for any correlation. Alternative methods of predicting the pathogenicity of the variants, such as M-CAP^26^, SIFT^27^, PolyPhen-2^28^, CADD^29^, and/or MetaLR^30^, could be used in conjunction with the metrics for the AF2 models to increase accuracy in predicting the mutational effects on function.

The metric, pLDDT, was introduced as a measure of the confidence in the AF2 models. In the context of obtaining alternate conformations, this metric is especially important for what appears to be a well folded structure. A recent report examining a curated subset of apo and ligand bound structures suggested that pLDDT values infer local conformational changes.^31^ The results presented here do not suggest that these regions of low confidence are incorrect in their two- or three-dimensional structure, but appear to be regions where there are contacts in the MSA that are not considered in the given structure (Fig. 2, Supp. Figs. 2, 3, 5, and 7). This is in agreement with a study that found that residues that had differential contacts in conformationally heterogenous proteins had multiple distance peaks in the distance matrix derived from MSAs.^32^ Therefore, these regions of uncertainty are targets for conformational switching as well as for direct contact points to consider for alternate conformations.

An expansion of this methodology would be to create a higher throughput work flow. This would allow for the ability to create a multitude of alternate conformations for both “native” and mutant proteins. One key element is the ability to determine the validity of the obtained structures as in some cases the mutations lead to unrealistic structures (Supp. Fig. 6D). This is where PCA can be applied to identify outliers in the structural landscape (Supplemental Information, Supp. Figs. 3 and 7).

The results presented here strongly support manipulation of the MSA to generate ensembles of multiple conformations of proteins via AF2. The PCA of the ensembles and conformations presented fully support the biochemical significance of the models generated here. These *in silico* structures can guide experimental design and be tested using spectroscopic approaches, and will ultimately provide a framework for interpretation of existing biochemical data and the development of mechanistic models.

## METHODS

### Initial Alphafold2 Structure

Protein sequences were downloaded from NCBI^33^. These sequences were input into the jupyter notebook for ColabFold.^6^ ColabFold implements folding of the protein with the model for Alphafold2 using MMseqs2 to generate the MSA. The notebook was used with the default parameters, which includes no template.^2,6,34–36^ The rationale for not including any templates is to allow Alphafold2 to generate structural intermediates that may not be achieved by the bias of including structural templates.

### In silico mutagensis

The choice of residues was based on previous information about the protein movement, either directly or from members in the family, the pLDDT values, and points of contact within the protein structure. At present, the method requires removal or alteration of contact points to alter the attention of Alphafold2. Once a region is chosen, all residue pairs of this region and the rest of the protein within 4 Å are tabulated. To keep from destabilizing secondary structure, any of the interacting partners that were within 4 amino acids in the linear sequence of the region of interest were omitted. Four angstroms was chosen as the cutoff to encompass polar and ionic interactions including those mediated by water. Alternate cutoffs or criteria could be chosen to yield smaller sets of potentially interacting pairs. Various methods of applying these pairs was carried out. In some cases, both sides of the pairs were mutated to allow complete disruption of the interaction at the region of interest. In other cases, only one side was chosen, either in the region of interest or in the residues outside this region allowing for a potentially new interaction to take place within the region of interaction.

### Modification of the MSA

The initial MSA generated by MMseqs2 is in a3m format. This format has the first sequence in all capitals without gaps. The subsequent sequences have a dash for a gap in that sequence and lowercase letters for a gap in the first sequence. The mutation across all sequences were made in the equivalent position without replacing gaps and were made to alanine unless otherwise noted in the Tables. The choice of alanine is to minimize any negative consequences of the mutation on secondary structure.

### Additional Alphafold2 Runs

The subsequent runs using the modified sequence and MSA were carried out with the ColabFold Alphafold2_advanced notebook with the following changes: the MSA method was set to single_sequence and the add_custom_msa checkbox was checked. This allowed for input of the modified MSA without any other MSA being used by Alphafold2.

### Additional Analysis

The residues used for the following analysis are given in the Tables. The TM score was carried out by TM_align.^37^ The RMSD calculation using 5 cycles of rejecting outliers, distance measurements, and figure generation were carried out in Pymol.^38^ The average TM and RMSD were calculated for the 10 combinations from a single Alphafold2 run. Principal component analysis was carried out with ProDy.^39^

## Acknowledgements

We thank Derek Claxton for helpful discussions and critically reading the manuscript. This work was supported by grants GM128087 and GM077659.

## Author Contributions

RAS: design, acquisition, and analysis. RAS and HSM: prepared the manuscript.

