## Supplemental Information for "Modeling Alternate Conformations with Alphafold2 via Modification of the Multiple Sequence Alignment"

**Additional Proteins:**

A recent report used an alternate methodology for sampling conformational space with Alphafold.[1] This method had to resort to using templates in one case, MCT1, and were unable to obtain the alternate conformation if one of the structures was present in the training set. To compare these two methods, four of the proteins from that study are examined here. Two of the proteins had no structures in the training set, LAT1 and MCT1, while two proteins MurJ and PfMATE only had one of the two conformations in the training set.

LAT1 is a human amino acid transporter belonging to the LeuT-fold family. The structure of LAT1 has been determined with an ancillary protein, 4f2hc, in multiple conditions. The structures compare here are inward open, 6irs,[2] and outward open, 7dsq, conformations (Supp. Fig. 12A). All the modeling was done in the absence of the ancillary protein. Running of the LAT1 sequence through ColabFold leads to an inward open structure (Supp. Fig. 12B, Supp. Table 7). Similar to Mhp1, TM5 was chosen as a region of interest. Mutation of residues within TM5 and interacting residues leads to models with movement in TMs 1a and 7 (Supp. Fig. 12C). Again, similar to Mhp1, residues with TM9/10 and interacting residues were for chosen. Mutation of these residue in the MSA leads to an altered conformation, with closing on the intracellular side and opening of the extracellular side (Supp. Fig. 12D). Based on the movement of TM1a, this region was chosen next for its effect on LAT1 models generated by Alphafold. Mutation of the MSA based on this region lead to a more diverse set of models (Supp. Fig. 12E). The plot of the TM-scores against the two structures indicate that our method is also able to obtain alternate conformations for LAT1 (Supp. Fig. 11).

MCT1 is an MFS protein that requires an ancillary protein, basigin, for expression and transport of monocarboxylates. The structure of MCT1 has been determined, in the presence of basigin, in an inward open conformation, 7da5, and an outward open conformation, 7ckr (Supp. Fig. 13A).[3] Similar to LAT1, all modeling was carried out in the absence of the ancillary protein. Input of the MCT1 sequence to ColabFold lead to the inward open structure (Supp. Fig. 13B). Similar to the other MFS proteins studied here three sets of mutations were carried, the whole interface between the two halves, the C-terminal half, or the N-terminal half (Supp. Table 13). Modification of the full interface leads to switching of the conformation to outward open (Supp. Fig. 13B, Supp. Table 8). Modification of either half of the interface leads to slight differences in the outward open conformation (Supp. Fig. 13C-D). For the N-terminal mutations, one model is misfolded with the two halves of the protein no longer in contact (Supp. Fig 13D). The plot of the TM-scores against the two structures indicate that this method is able to obtain alternate conformations for MCT1 without a template structure to bias Alphafold2 (Supp. Fig. 11).

MuJ is an *Escherichia coli* Lipid II flippase. The structure has been determined in an inward open, 5t77,[4] and outward open, 6nc9,[5] conformation (Supp. Fig. 14A). Input of the protein sequence to ColabFold generates five models that are inward open (Supp. Fig. 14B). Similar to the MFS proteins, MurJ was split in half and the interaction region between the two halves was explored (Supp. Table 9). This choice leads to an alteration of conformation that by TM-score approaches the outward open structure (Supp. Table 9). Examination of the structures indicate that TM13 and TM14 have a different conformation compared to the experimentally determined structure (Supp. Fig. 14C). The pLDDT values for the initial run of MurJ indicate some uncertainty in the structure that corresponds to the last helix in the N-terminal portion of the flippase. The next three modifications of the MSA focus on this region and the loop connecting the two halves of the protein. These modifications lead to a variety in the extent of opening of the protein (Supp. Fig. 14D-F). The TM-score plot does show that the method is able to obtain models comparable to the experimentally obtained structure (Supp. Fig. 11). The cluster of models around 0.7 on the x-axis are for the models where the position of TM13 and TM14 are different than in the experimental structure. This highlights the need to manually inspect the resultant models or carry out additional analysis as distortions might not be readily apparent from the TM scores.

PfMATE is a multidrug and toxic compound extrusion (MATE) protein *Pyrococcus furiosus.* The structure for PfMATE has been obtained for both an inward open, 6fhz[6], and outward open, 3vvn[7] (Supp. Fig. 15A). Input of the protein sequence for PfMATE to ColabFold generates five models that are outward open (Supp. Fig. 15B). Similar to the MFS proteins, PfMATE was split in half and the interaction region between the two halves was explored (Supp. Table 10). The Alphafold2 models for this modification of the MSA and PfMATE sequence leads to the alternate inward open conformation (Supp. Fig. 15C). Models generated by using only the C- or N-terminal halves of the protein also lead to inward open conformations, though there are differences between the three choices (Supp. Fig. 15D-E). The TM-score plot does not exhibit a broad array of structures as all of the additional models cluster near one another (Supp. Fig. 11). While intermediates might not have been generated, it is clear that this methodology is able to obtain the alternate conformation.

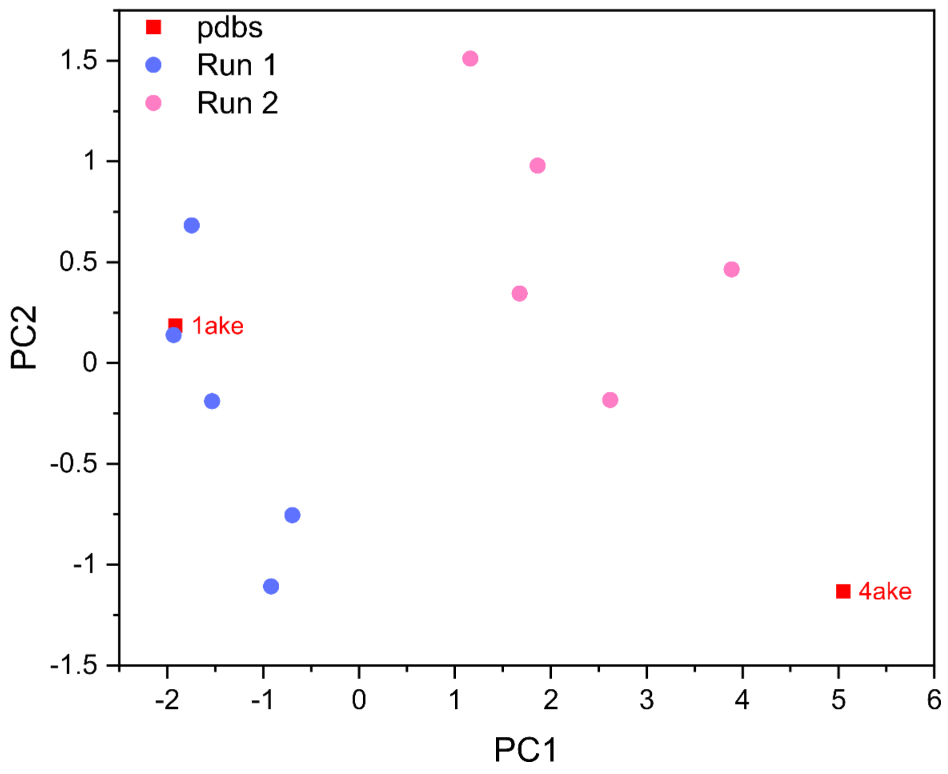

Supplemental Figure 1: Adenylate Kinase Principal Component Analysis. Plot of the first two components against each other. The two Alphafold2 runs appear to segregate into two different conformations.

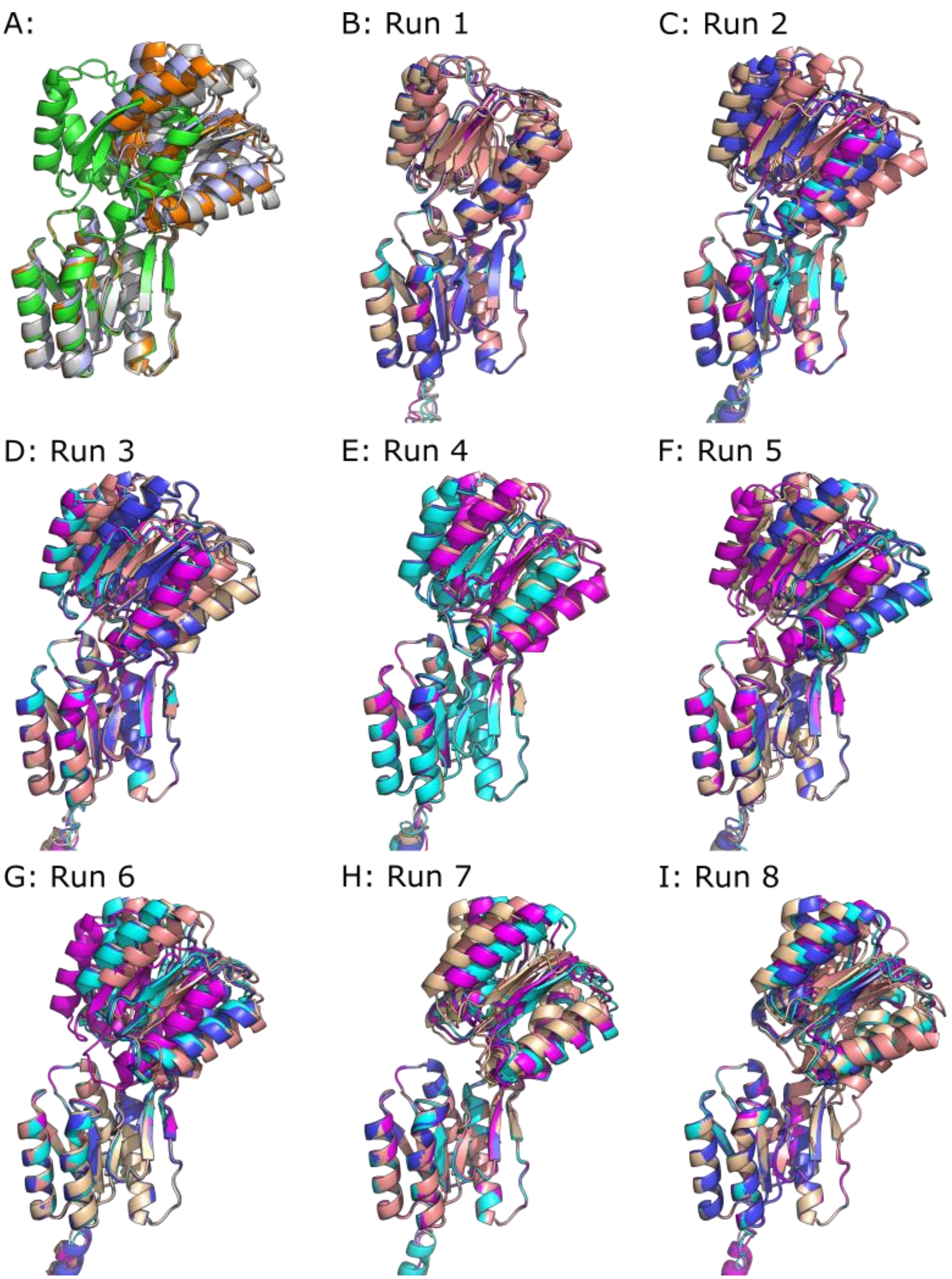

Supplemental Figure 2: Structures of E. coli ribose binding protein. For display the models were aligned with the crystal structures using residues 1-40 in Pymol. A) Four crystal structures: 2dri (green), 1urp (light blue), 1ba2B (orange), and 1ba2A (grey). B-I) The five Alphafold2 models for Runs 1-8, respectively.

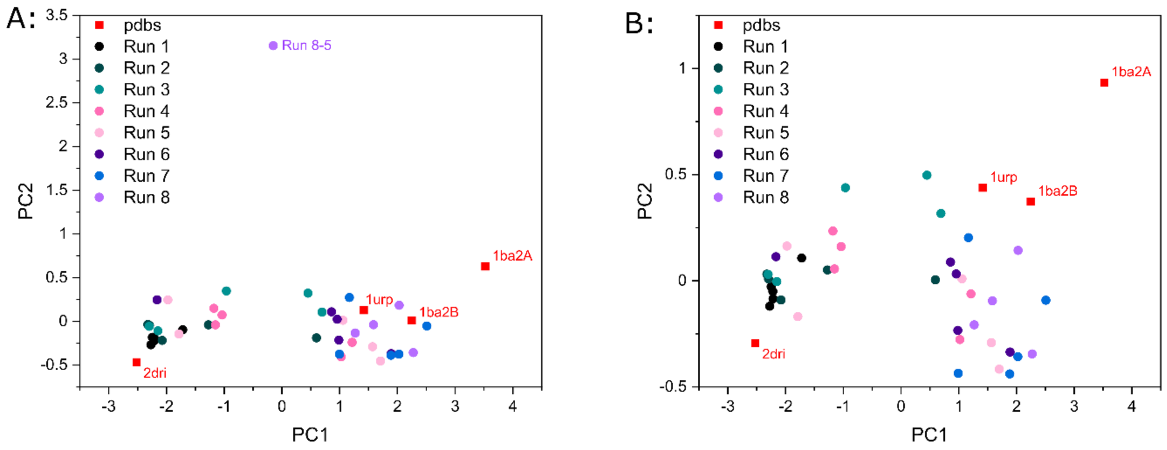

Supplemental Figure 3: Ribose Binding Protein Principal Component Analysis. A) Initial PC analysis for all structures. B) Reanalysis removing the apparent outlier, Run 8-5.

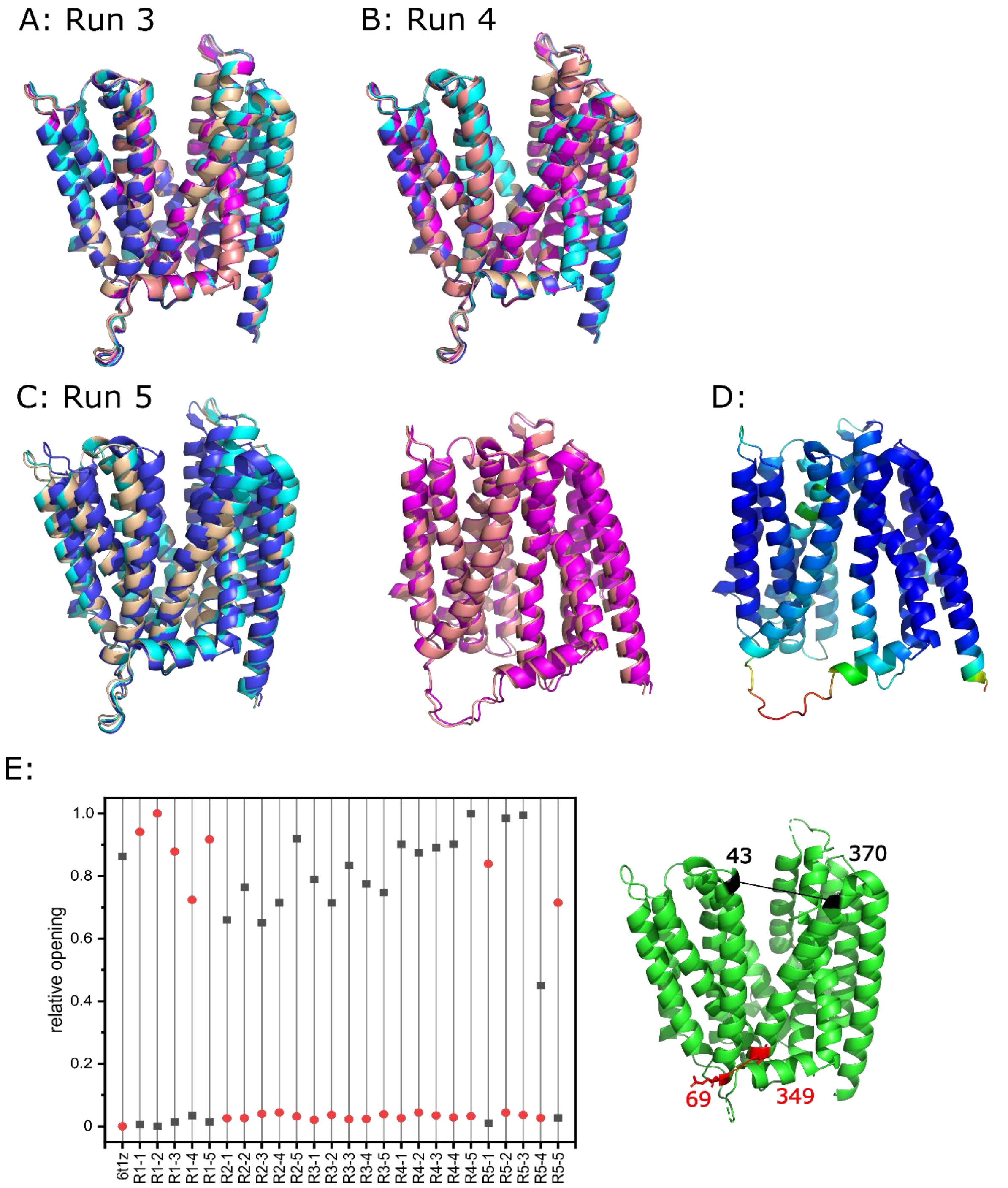

Supplemental Figure 4: Additional structures for LmrP. A) Five Alphafold2 models after mutation of the N-terminal residues at the interface. B) Five Alphafold2 models after mutation of the C-terminal residues at the interface. C) Effect of mutation of the 3 residues at the middle of the interface. On the left are 3 outward facing models and on the right are the 2 inward facing models. D) Plot of model 1 from Run 1 colored by pLDDT. E) Plot of the extent of opening of the intracellular (red, distance between residues 69 and 349) and extracellular (black, distance between 43 and 370) vestibule.

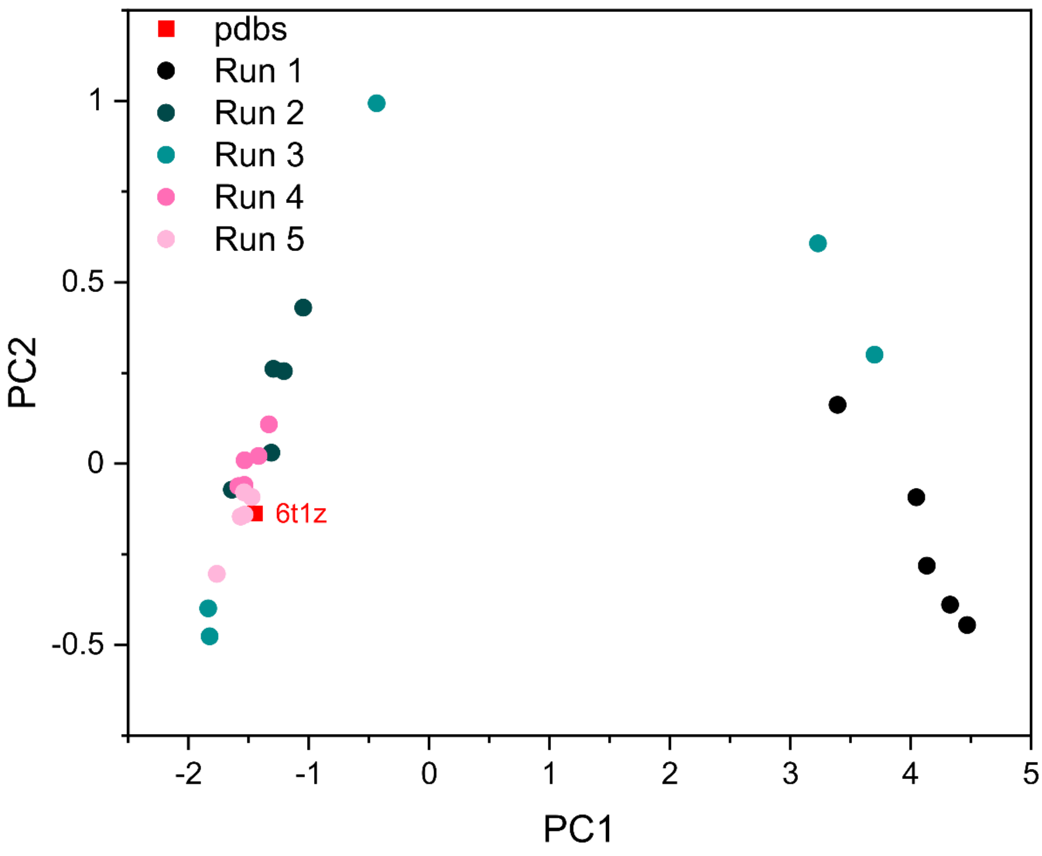

Supplemental Figure 5: LmrP Principal Component Analysis. Plot of the first two components relative to each other.

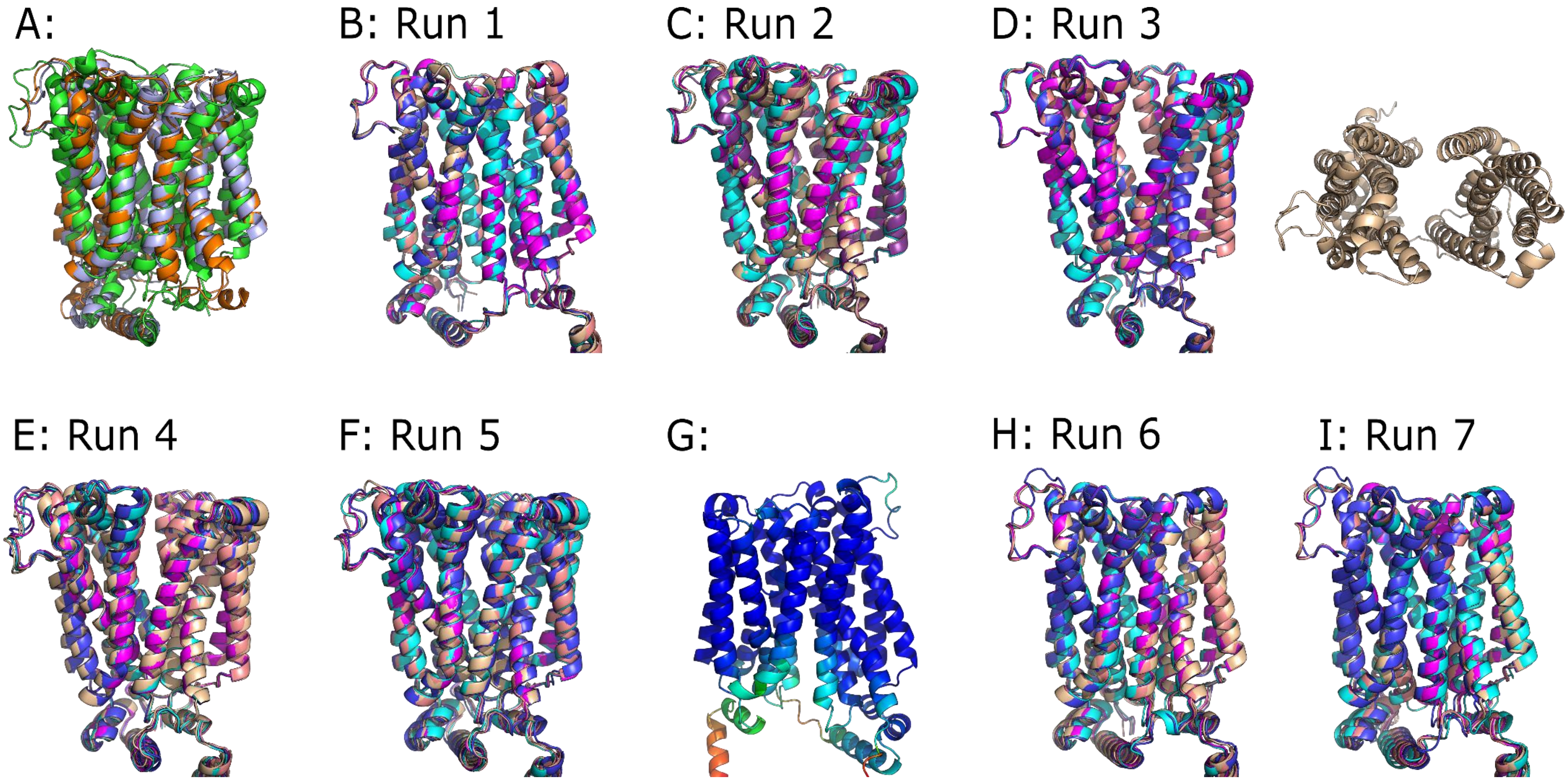
A:
B: Run 1
C: Run 2
D: Run 3
E: Run 4
F: Run 5
G:
H: Run 6
I: Run 7

J:

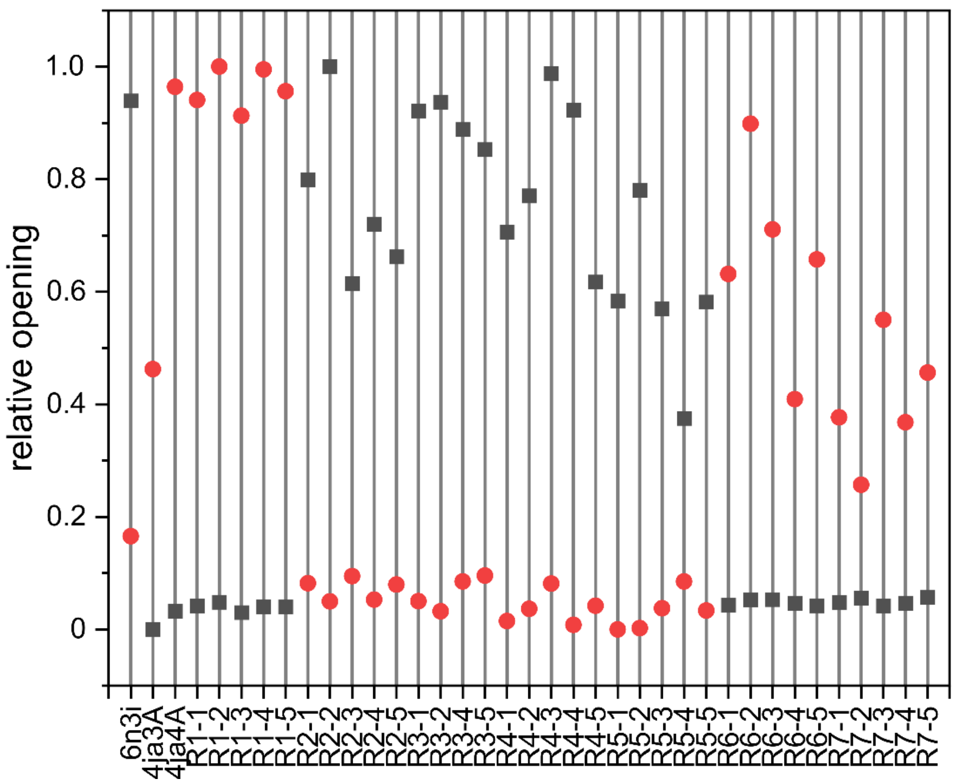

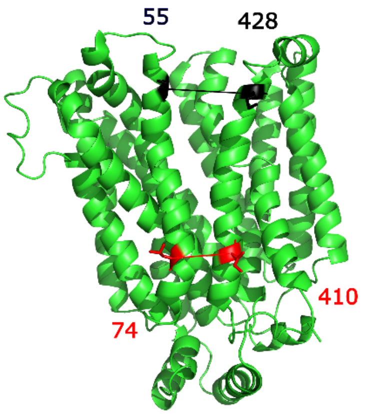

Supplemental Figure 6: Structures of E. coli MFS symporter XylE. A) Three crystal structures: 6n3i (green), 4ja3 (light blue), and 4ja4 (orange). B) The five Alphafold2 models for the input protein sequence. C) Five Alphafold2 models after mutation of residues at the interface of the inward facing structure from the top model in B. D) Alphafold2 models after mutation of the C-terminal residues at the interface. On the left are the four models that match the outward open structure, 6n3i. On the right is the top view of the one models where the two halves are no longer interacting (third ranked model by pLDDT). E) Five Alphafold2 models after mutation of the N-terminal residues at the interface. F) Five Alphafold2 models for the MSA with the double mutation G58W/L315W. G) The top model from A rotated 180° and colored by pLDDT. The region at the bottom of the two facing helices were used for Run 6 and Run 7. H) Five Alphafold2 models for the MSA with mutation of residues and their contacts in the region 154-173. I) Five Alphafold2 models for the MSA with mutation of residues and their contacts in the region 327-341. J) Plot of the extent of opening of the intracellular (red) and extracellular (black) vestibules.

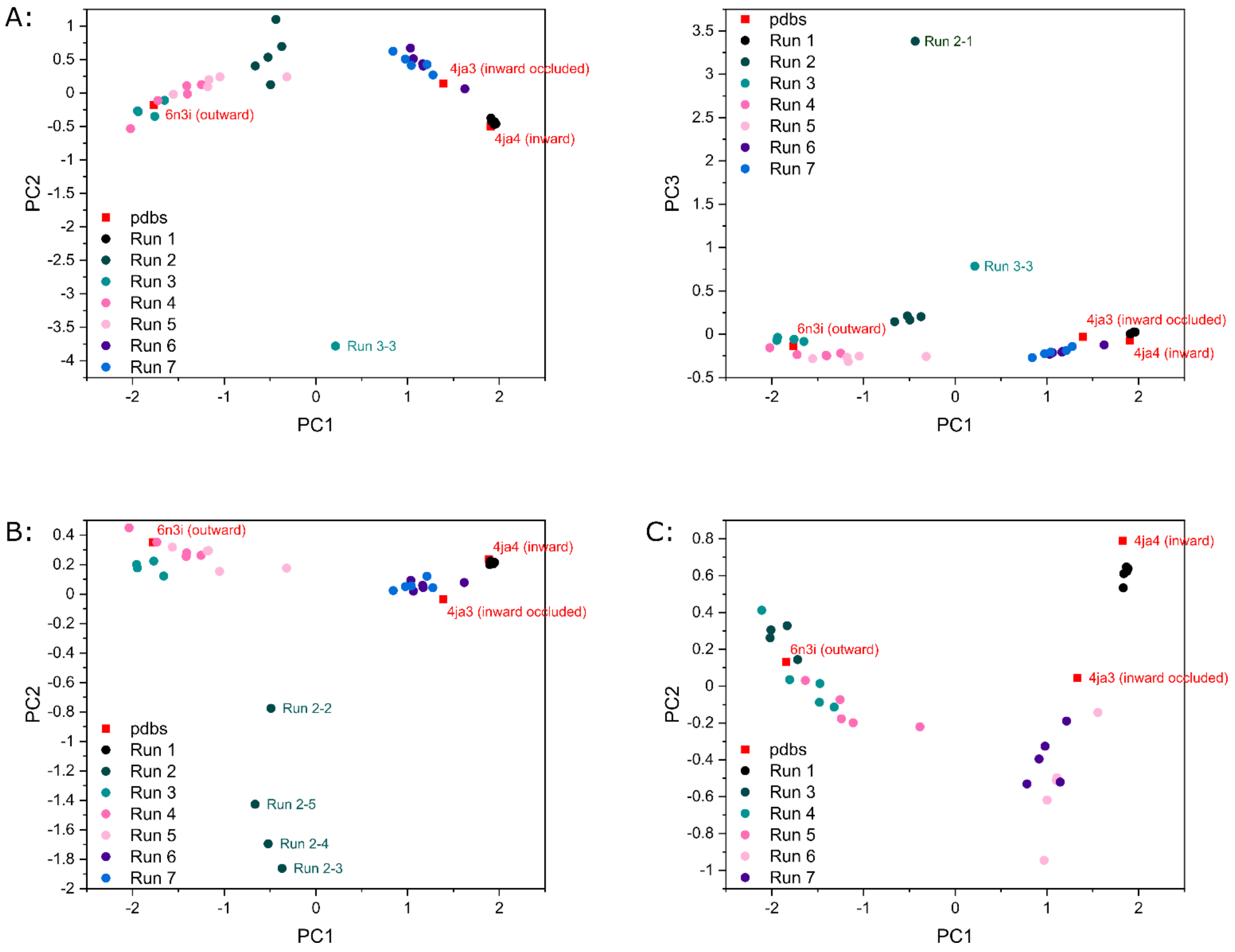

Supplemental Figure 7: XylE Principal Component Analysis: A) Analysis using all structures and models. Principal component 1 plotted vs component 2 (left) and component 3 (right). B) Plot of the first component vs the second component for the analysis removing Run 2-1 and 3-3 models. C) Plot of the first component vs the second component for the analysis removing the rest of Run 2.

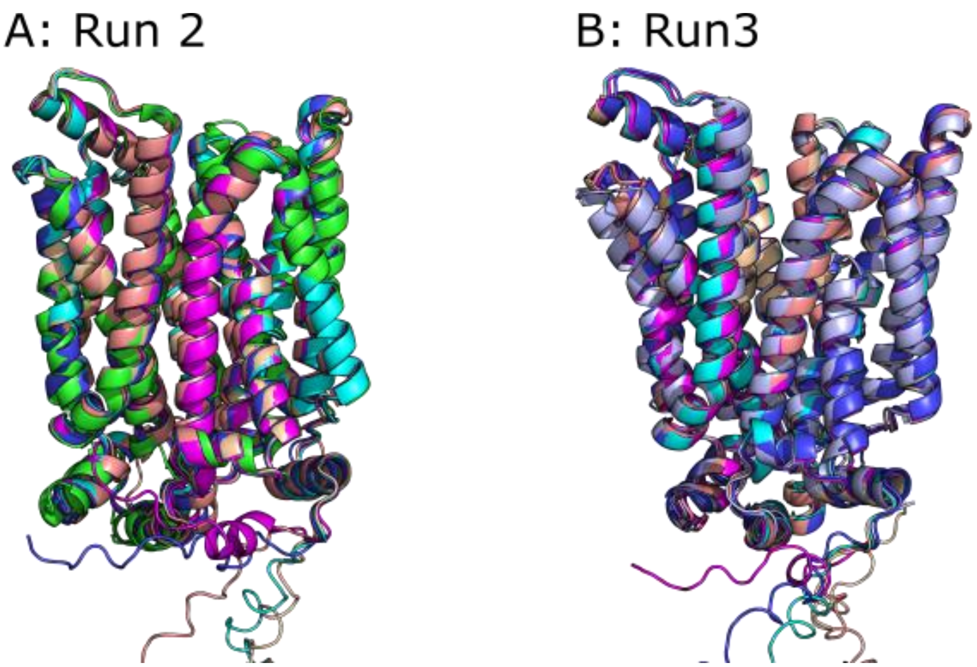

Supplemental Figure 8: Structures of GLUT5. A) Five Alphafold2 models superimposed with the inward open structure, 4yb9 (green) . B) Five Alphafold2 models superimposed with the outward open structure, 4ybq (light blue).

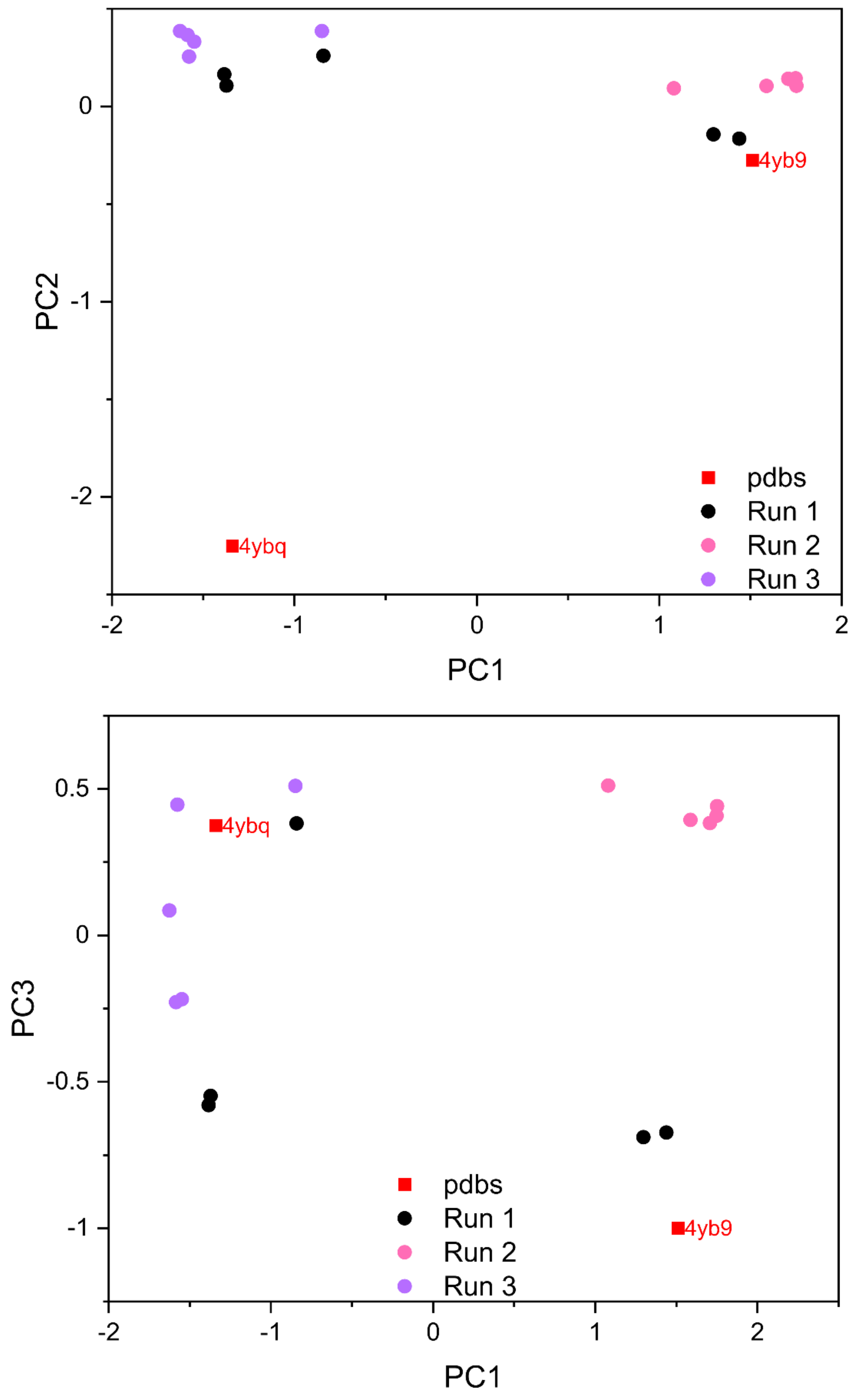

Supplemental Figure 9: GLUT5 Principal Component Analysis. Plot of the first component vs the second component (top) and third component (bottom).

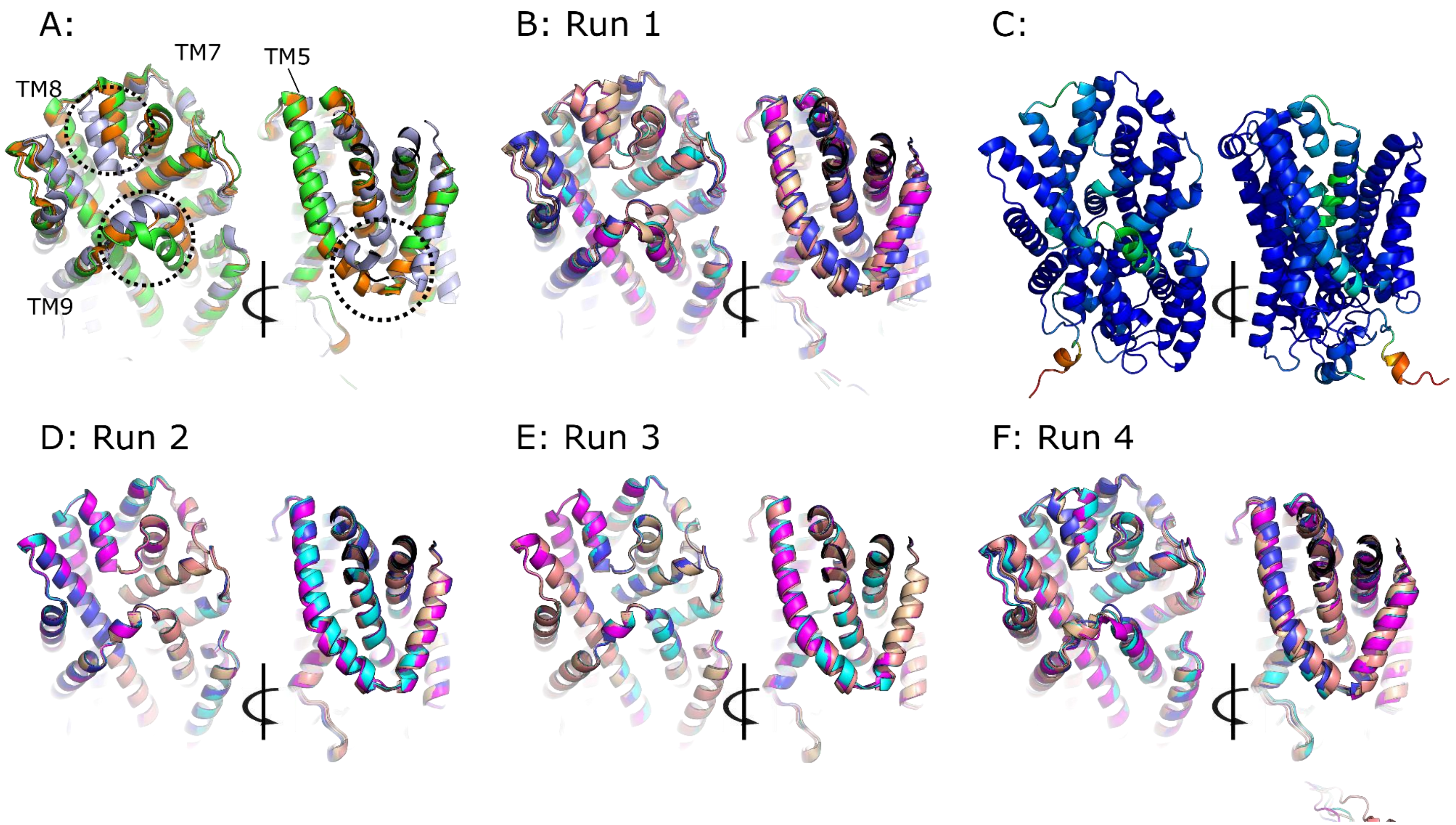

Supplemental Figure 10: Structures of MHP1. On the left is a view at the extracellular vestibule and on the right the protein is rotated ~180°. A) Three conformations obtained from x-ray crystallography, 2jln (green), 2x79 (grey), and 4d1b (orange). Note the differences in the loop between helices 7 and 8 (top circle), helices 9-10 (bottom circle) in the outer vestibule, and in helix 5. B) Five Alphafold2 models that catch the various conformations in the three structures in A. C) The top model from B colored by LDDT score. Note that the less certain structural elements correspond to the regions with different conformations in the crystal structures. D) Five Alphafold2 models based on the interaction region 159-170. E) Five Alphafold2 models based on the interaction region 159-170 choosing only the sites opposite this region. F) Five Alphafold2 models based on the interaction region 352-368.

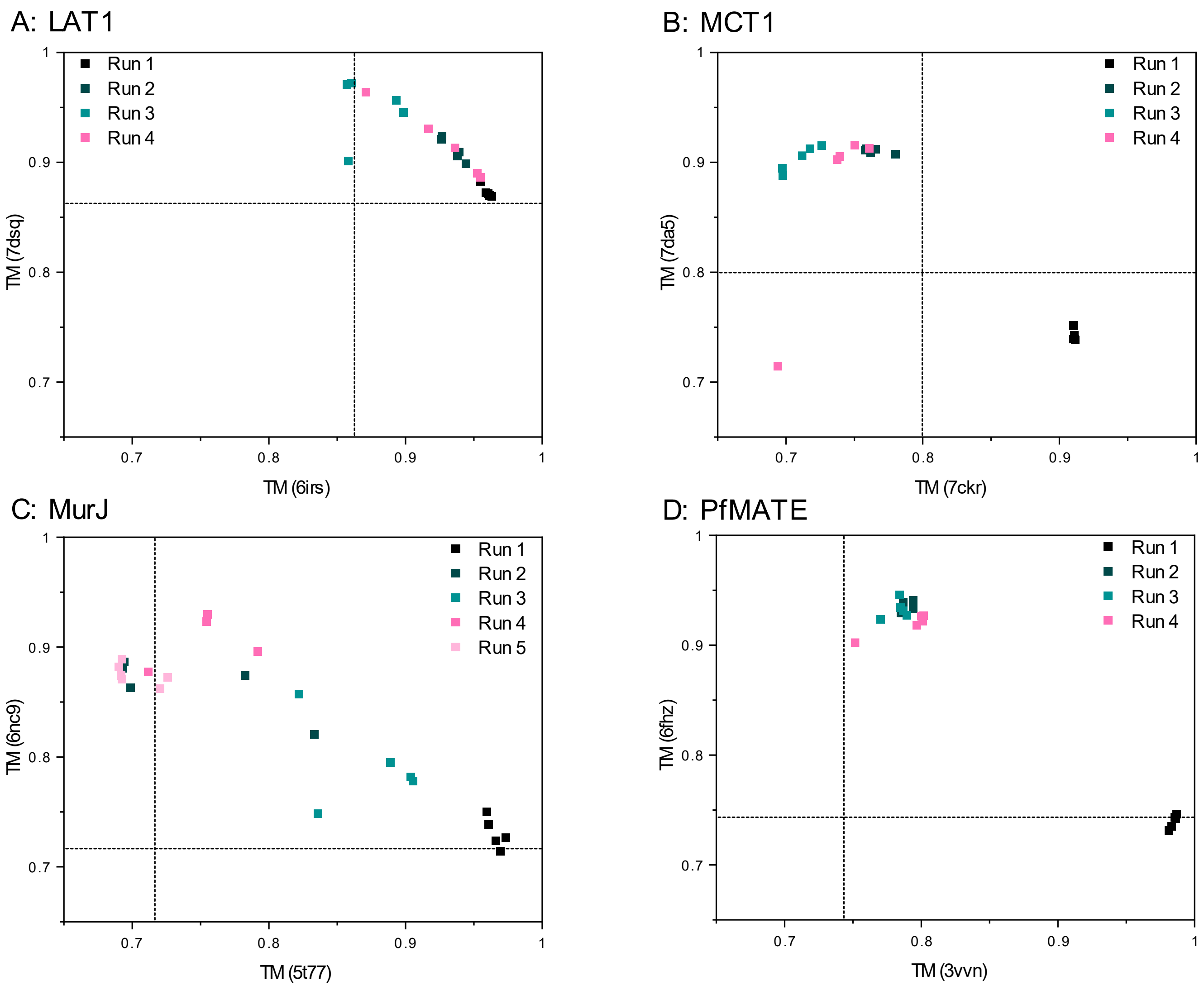

Supplemental Figure 11: TM-scores of the models relative to the crystal structures plotted against each other. The dashed line in each plot is the TM-score between the two crystal structures. A) LAT1. B) MCT1. C) MurJ. D) PfMATE.

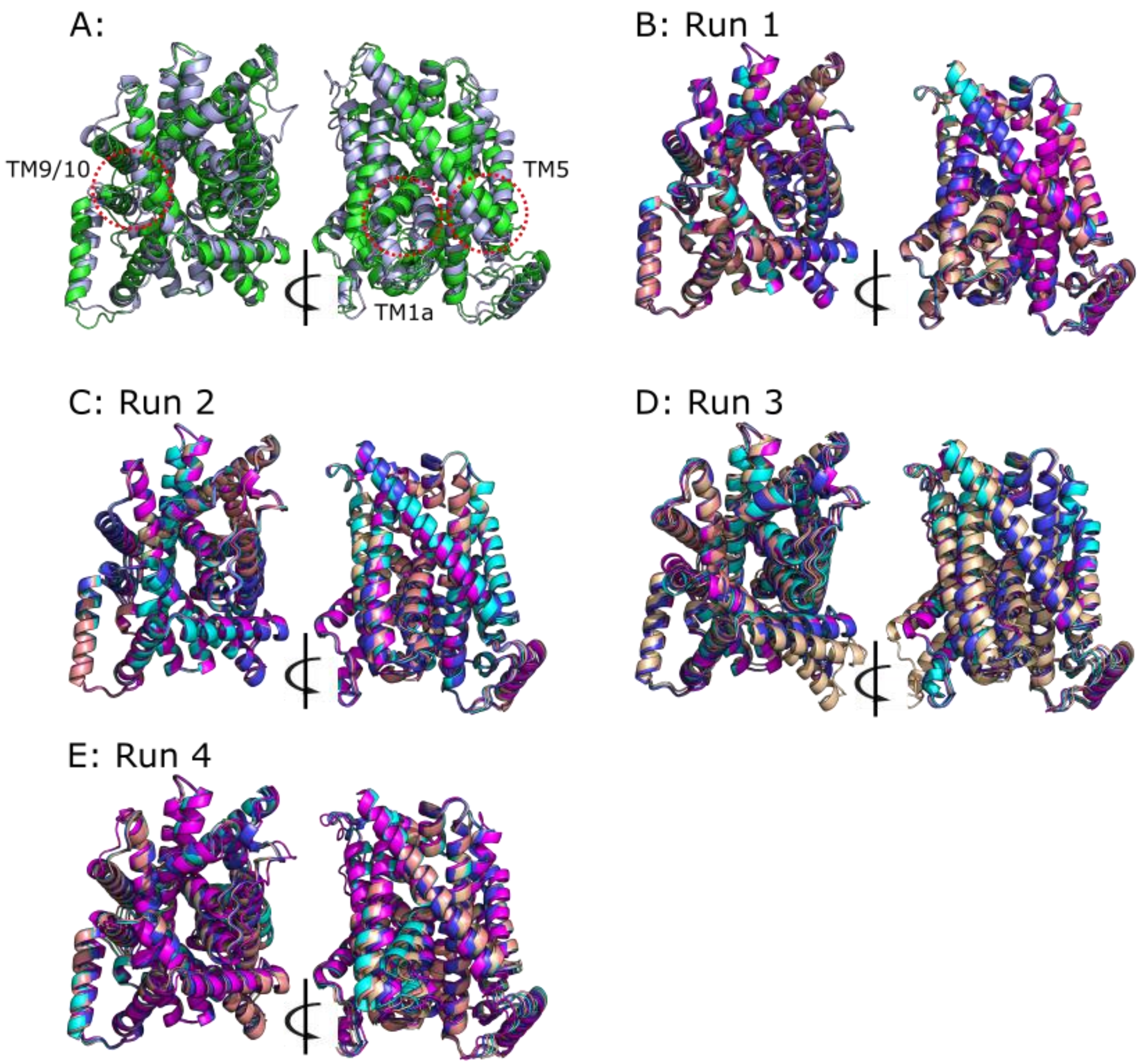

Supplemental Figure 12: Structures of MHP1. On the left is a view at the extracellular vestibule and on the right the protein is rotated ~180°. The first 50 residues were removed for visualization. A) PDB structures 6irs (green) and 7dsq (light blue). B) Alphafold2 models for the default MSA. C) Modification based on TM5. D) Modification based ont TM 9/10. E) Modification based on TM1a.

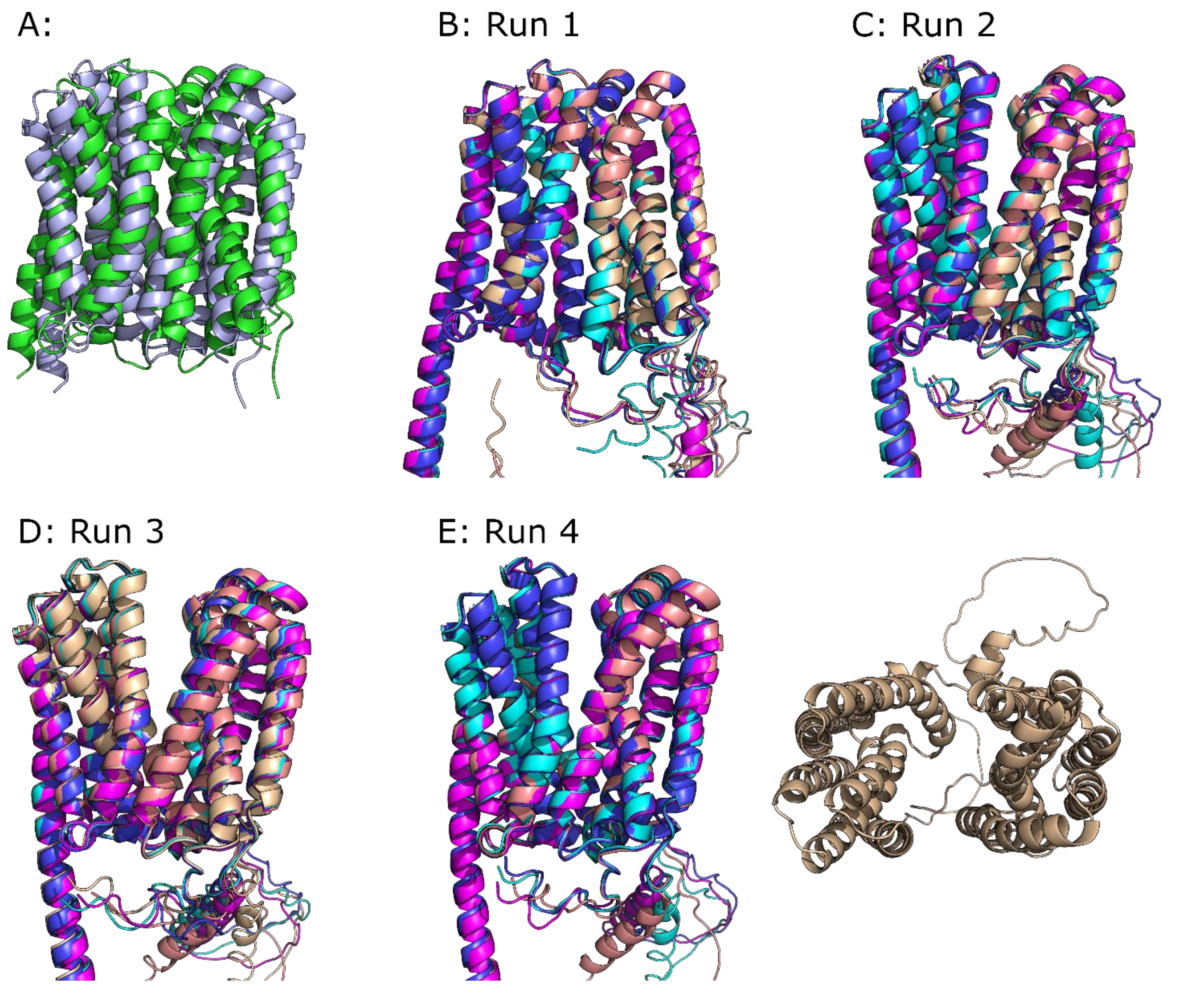

Supplemental Figure 13: Structures of MCT1. A) PDB structures 7da5 (green) and 7ckr (light blue). B) Five Alphafold2 models from the defualt run. C) Five Alphafold2 models after mutation of the residues at the interface. D) Five Alphafold2 models after mutation of the C-terminal residues at the interface. E) Five Alphafold2 models after mutation of the N-terminal residues at the interface.

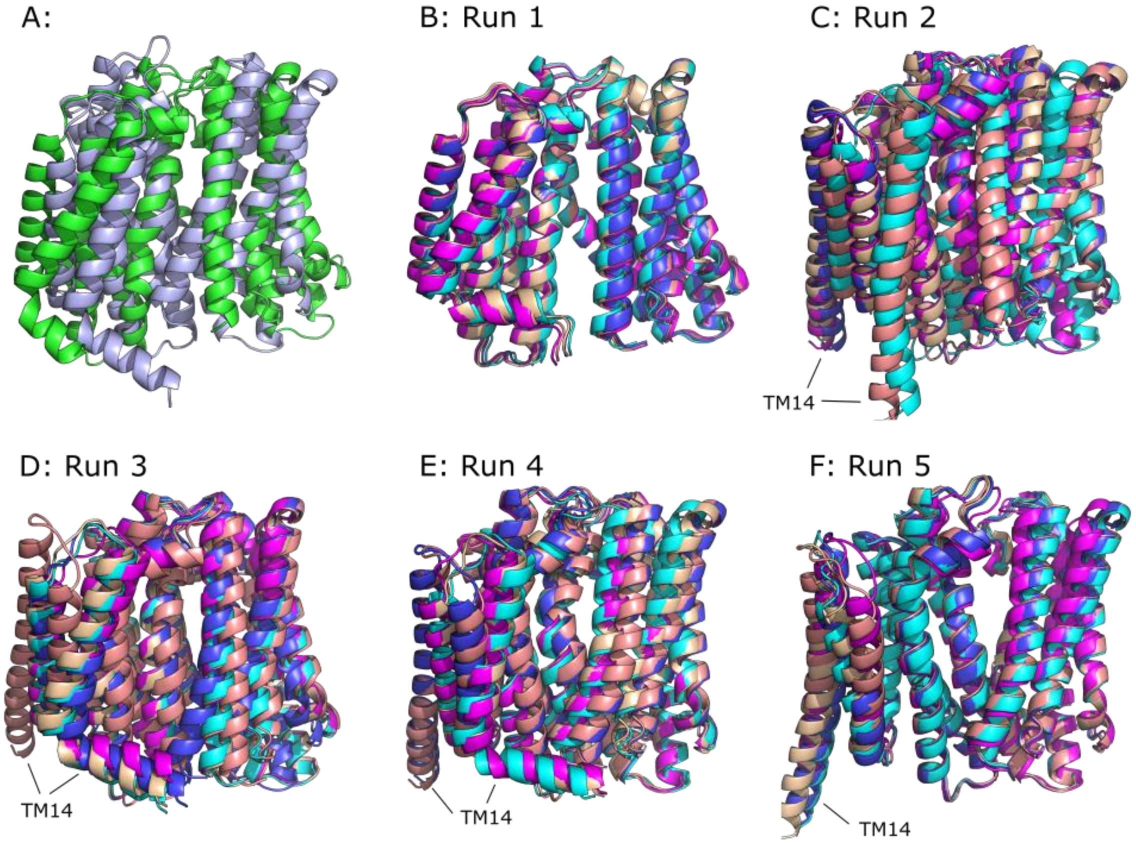

Supplemental Figure 14: Structures of MurJ. A) PDB structures 5t77 (green) and 6nc9 (light blue). B) Five Alphafold2 models from the default run. C) Five Alphafold2 models after mutation of the residues at the interface. D) Five Alphafold2 models using residues 223-234. E) Five Alphafold2 models using residues 248-255. F) Five Alphafold2 models using residues 230-241.

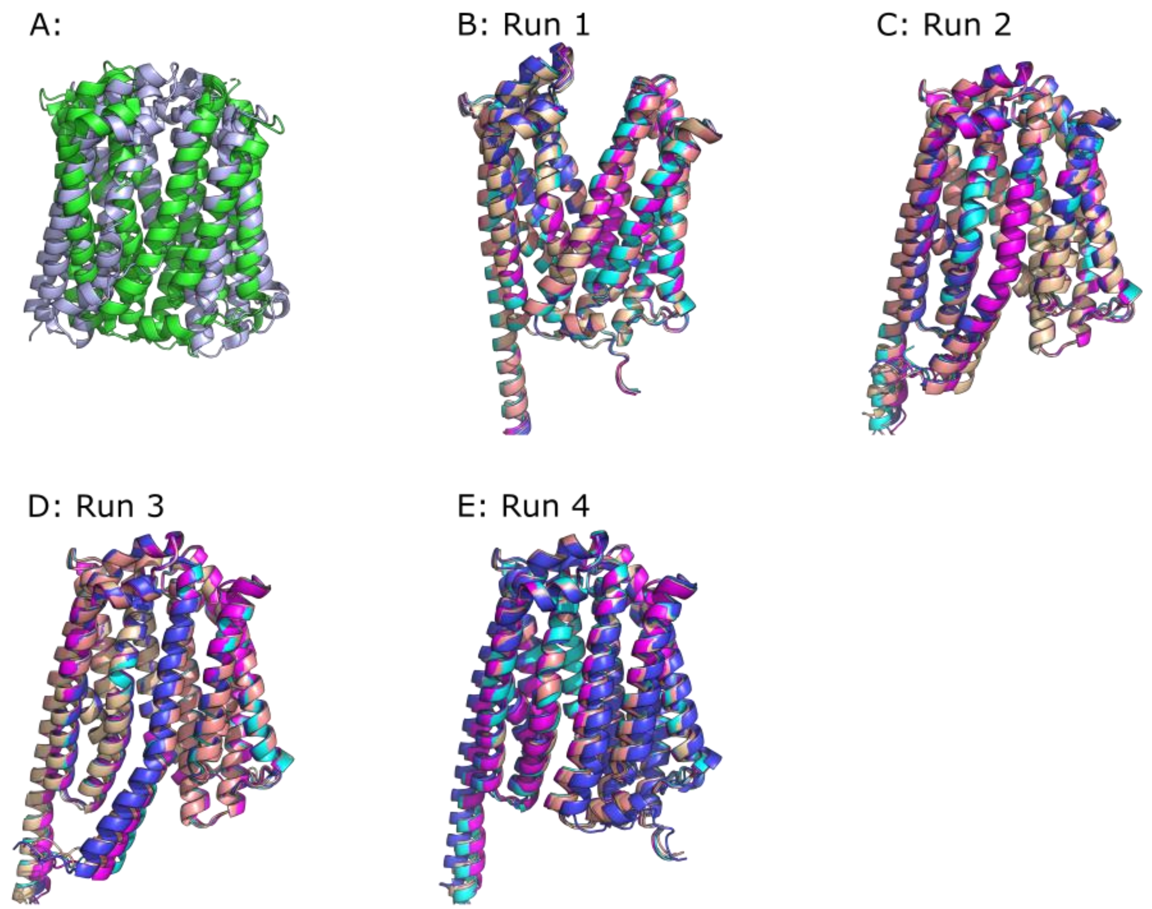

Supplemental Figure 15: Structures of PfMATE. A) PDB structures 3vvn (green) and 6fhz (light blue). B) Five Alphafold2 models from the default run. C) Five Alphafold2 models after mutation of the residues at the interface. D) Five Alphafold2 models after mutation of the C-terminal residues at the interface. E) Five Alphafold2 models after mutation of the N-terminal residues at the interface.

| Supplemental Table 1: Adenylate Kinase | | | | | | | | | | | | | |
|---|---|---|---|---|---|---|---|---|---|---|---|---|---|
| | Average | | | Model 1 | | Model 2 | | Model 3 | | Model 4 | | Model 5 | |
| Run | TM | RMSD | PDB | TM | RMSD | TM | RMSD | TM | RMSD | TM | RMSD | TM | RMSD |
| 1 | 0.956 | 0.362 | 1ake | 0.98 | 0.56 | 0.92 | 0.59 | 0.97 | 0.58 | 0.97 | 0.75 | 0.93 | 0.65 |
| | | | 4ake | 0.69 | 1.93 | 0.71 | 2.00 | 0.70 | 1.97 | 0.71 | 1.66 | 0.72 | 1.96 |
| 2 | 0.920 | 0.626 | 1ake | 0.70 | 2.23 | 0.79 | 2.00 | 0.77 | 1.73 | 0.83 | 1.91 | 0.80 | 1.90 |
| | | | 4ake | 0.88 | 1.72 | 0.81 | 1.46 | 0.84 | 1.64 | 0.78 | 1.48 | 0.80 | 1.45 |

| Protein Accession Number/Region: Mutated Residues | |
|---|---|
| 1 | WP_001220233 |
| 2 | Region 114-173: 7, 8, 9, 10, 11, 33, 36, 54, 55, 56, 58, 115, 116, 119, 120, 156, 163, 166, 167, 168, 170, 171, 198, 199 |

The TM and RMSD were calculated using residues 1-214.

The average TM and RMSD are the average values for the five models against each other in a single run (10 total comparisons).

Numbers in blue are the best TM or RMSD score for that structure.

The residues were mutated to Alanine.

| Supplemental Table 2: Ribose Binding Protein | | | | | | | | | | | | | |
|---|---|---|---|---|---|---|---|---|---|---|---|---|---|
| | Average | | | Model 1 | | Model 2 | | Model 3 | | Model 4 | | Model 5 | |
| Run | TM | RMSD | PDB | TM | RMSD | TM | RMSD | TM | RMSD | TM | RMSD | TM | RMSD |
| 1 | 0.951 | 0.396 | 1ba2A | 0.61 | 6.05 | 0.61 | 6.05 | 0.60 | 6.12 | 0.60 | 6.05 | 0.62 | 5.55 |
| | | | 1ba2B | 0.69 | 4.64 | 0.68 | 4.63 | 0.68 | 4.70 | 0.68 | 4.64 | 0.70 | 4.10 |
| | | | 1urp | 0.73 | 3.82 | 0.73 | 3.83 | 0.72 | 3.89 | 0.73 | 3.83 | 0.75 | 3.28 |
| | | | 2dri | 0.90 | 0.56 | 0.90 | 0.52 | 0.90 | 0.47 | 0.90 | 0.52 | 0.88 | 1.15 |
| 2 | 0.895 | 1.489 | 1ba2A | 0.62 | 6.08 | 0.61 | 6.10 | 0.62 | 5.90 | 0.66 | 5.08 | 0.75 | 3.28 |
| | | | 1ba2B | 0.68 | 4.70 | 0.68 | 4.73 | 0.69 | 4.48 | 0.73 | 3.69 | 0.84 | 1.81 |
| | | | 1urp | 0.73 | 3.90 | 0.73 | 3.93 | 0.74 | 3.39 | 0.77 | 2.91 | 0.88 | 1.09 |
| | | | 2dri | 0.90 | 0.69 | 0.90 | 0.65 | 0.90 | 0.70 | 0.86 | 1.45 | 0.76 | 3.29 |
| 3 | 0.875 | 1.971 | 1ba2A | 0.62 | 6.09 | 0.62 | 5.95 | 0.74 | 3.32 | 0.76 | 3.09 | 0.67 | 4.72 |
| | | | 1ba2B | 0.68 | 4.70 | 0.69 | 4.54 | 0.83 | 1.95 | 0.85 | 1.66 | 0.75 | 3.426 |
| | | | 1urp | 0.73 | 3.91 | 0.74 | 3.75 | 0.88 | 1.19 | 0.89 | 0.93 | 0.78 | 2.69 |
| | | | 2dri | 0.90 | 0.64 | 0.90 | 0.70 | 0.76 | 3.25 | 0.75 | 3.45 | 0.83 | 1.97 |
| 4 | 0.880 | 1.565 | 1ba2A | 0.76 | 2.99 | 0.66 | 4.99 | 0.66 | 4.98 | 0.67 | 4.85 | 0.78 | 2.73 |
| | | | 1ba2B | 0.86 | 1.59 | 0.74 | 3.63 | 0.74 | 3.65 | 0.75 | 3.52 | 0.87 | 1.29 |
| | | | 1urp | 0.88 | 1.08 | 0.77 | 2.85 | 0.77 | 2.92 | 0.78 | 2.79 | 0.89 | 0.82 |
| | | | 2dri | 0.73 | 3.70 | 0.85 | 1.63 | 0.85 | 1.72 | 0.84 | 1.81 | 0.72 | 3.92 |
| 5 | 0.833 | 2.353 | 1ba2A | 0.62 | 5.66 | 0.79 | 2.49 | 0.62 | 5.79 | 0.77 | 2.83 | 0.79 | 2.46 |
| | | | 1ba2B | 0.70 | 4.22 | 0.88 | 1.15 | 0.69 | 4.43 | 0.86 | 1.45 | 0.88 | 1.11 |
| | | | 1urp | 0.75 | 3.43 | 0.88 | 0.84 | 0.74 | 3.60 | 0.89 | 0.79 | 0.88 | 0.98 |
| | | | 2dri | 0.88 | 1.03 | 0.69 | 4.29 | 0.88 | 1.08 | 0.72 | 3.79 | 0.70 | 4.42 |
| 6 | 0.885 | 1.900 | 1ba2A | 0.60 | 6.00 | 0.76 | 2.96 | 0.76 | 2.89 | 0.76 | 2.97 | 0.81 | 2.25 |
| | | | 1ba2B | 0.68 | 4.64 | 0.85 | 1.64 | 0.86 | 1.54 | 0.85 | 1.66 | 0.88 | 0.95 |
| | | | 1urp | 0.73 | 3.83 | 0.88 | 1.13 | 0.89 | 0.93 | 0.88 | 1.04 | 0.88 | 1.02 |
| | | | 2dri | 0.88 | 0.96 | 0.72 | 3.68 | 0.73 | 3.69 | 0.73 | 3.60 | 0.68 | 4.60 |
| 7 | 0.943 | 0.990 | 1ba2A | 0.80 | 2.30 | 0.85 | 1.60 | 0.76 | 3.05 | 0.82 | 2.14 | 0.78 | 2.66 |
| | | | 1ba2B | 0.88 | 1.01 | 0.88 | 0.70 | 0.85 | 1.71 | 0.88 | 0.86 | 0.86 | 1.47 |
| | | | 1urp | 0.87 | 1.02 | 0.86 | 1.37 | 0.87 | 1.22 | 0.87 | 1.06 | 0.88 | 0.94 |
| | | | 2dri | 0.68 | 4.63 | 0.64 | 5.26 | 0.72 | 3.69 | 0.67 | 4.76 | 0.72 | 3.94 |
| 8 | 0.872 | 2.090 | 1ba2A | 0.84 | 1.89 | 0.80 | 2.36 | 0.78 | 2.70 | 0.83 | 1.95 | 0.68 | 4.52 |
| | | | 1ba2B | 0.89 | 0.75 | 0.88 | 1.10 | 0.86 | 1.43 | 0.88 | 0.80 | 0.70 | 4.00 |
| | | | 1urp | 0.88 | 0.98 | 0.89 | 0.73 | 0.88 | 0.94 | 0.86 | 1.26 | 0.72 | 3.53 |
| | | | 2dri | 0.67 | 4.80 | 0.70 | 4.31 | 0.71 | 3.99 | 0.66 | 5.01 | 0.67 | 4.48 |

| Protein Accession Number/Region: Mutated Residues | |
|---|---|
| 1 | WP_001056273 |
| 2 | D67R |
| 3 | Region 155-161: 91, 92, 93, 153, 155, 159, 160, 161 |
| 4 | Run 2 + Run 3 |
| 5 | Regions 127-130, 259-262, 287-291: 40, 114, 126, 127, 240, 257, 259, 260, 262, 288, 291 |
| 6 | Run 2 + Run 5 |
| 7 | Run 3 + Run 5 |
| 8 | Run 2 + Run 7 |

The TM and RMSD were calculated using residues 1-271.
The average TM and RMSD are the average values for the five models against each other in a single run (10 total comparisons).
Numbers in blue are the best TM or RMSD score for that structure.
Numbers in red are the best TM or RMSD score for that structure, but not for that model indicating that this structure may not be represented in the models.
The residues were mutated to Alanine.

| Supplemental Table 3: LmrP | | | | | | | | | | | | | |
|---|---|---|---|---|---|---|---|---|---|---|---|---|---|
| | Average | | | Model 1 | | Model 2 | | Model 3 | | Model 4 | | Model 5 | |
| Run | TM | RMSD | PDB | TM | RMSD | TM | RMSD | TM | RMSD | TM | RMSD | TM | RMSD |
| 1 | 0.984 | 0.775 | 6t1z | 0.69 | 5.89 | 0.69 | 5.97 | 0.71 | 5.62 | 0.73 | 4.99 | 0.70 | 5.69 |
| 2 | 0.988 | 0.409 | 6t1z | 0.96 | 0.85 | 0.96 | 0.81 | 0.96 | 0.99 | 0.96 | 0.85 | 0.96 | 0.80 |
| 3 | 0.998 | 0.267 | 6t1z | 0.97 | 0.75 | 0.97 | 0.78 | 0.97 | 0.70 | 0.97 | 0.74 | 0.97 | 0.78 |
| 4 | 0.996 | 0.290 | 6t1z | 0.97 | 0.67 | 0.97 | 0.68 | 0.97 | 0.69 | 0.97 | 0.68 | 0.97 | 0.74 |
| 5 | 0.830 | 3.377 | 6t1z | 0.73 | 5.22 | 0.97 | 0.79 | 0.97 | 0.76 | 0.94 | 1.55 | 0.75 | 4.80 |

| Protein Accession Number/Region: Mutated Residues | |
|---|---|
| 1 | WP_011836081 |
| 2 | N/C split (186/215): 29, 33, 34, 37, 42, 43, 46, 47, 49, 53, 54, 57, 151, 154, 155, 158, 159, 162, 235, 236, 240, 244, 257, 259, 260, 263, 270, 359, 362, 363, 366, 369, 370, 373 |
| 3 | N/C split (186/215) N-term only: 29, 33, 34, 37, 42, 43, 46, 47, 49, 53, 54, 57, 151, 154, 155, 158, 159, 162 |
| 4 | N/C split (186/215) C-term only: 235, 236, 240, 244, 257, 259, 260, 263, 270, 359, 362, 363, 366, 369, 370, 373 |
| 5 | C270A, P359A, I363A |

The TM and RMSD were calculated using residues 1-406.

The average TM and RMSD are the average values for the five models against each other in a single run (10 total comparisons).

Numbers in blue are the best TM or RMSD score for that structure.

The residues were mutated to Alanine unless otherwise noted.

Supplemental Table 4: XylE

| | Average | | | Model 1 | | Model 2 | | Model 3 | | Model 4 | | Model 5 | |
|---|---|---|---|---|---|---|---|---|---|---|---|---|---|
| Run | TM | RMSD | PDB | TM | RMSD | TM | RMSD | TM | RMSD | TM | RMSD | TM | RMSD |
| 1 | 0.998 | 0.177 | 4ja3 | 0.94 | 0.63 | 0.94 | 0.66 | 0.94 | 0.66 | 0.94 | 0.69 | 0.94 | 0.60 |
| | | | 4ja4 | 0.95 | 0.78 | 0.95 | 0.79 | 0.95 | 0.80 | 0.95 | 0.80 | 0.95 | 0.78 |
| | | | 6n3i | 0.82 | 4.02 | 0.83 | 4.06 | 0.82 | 4.07 | 0.82 | 4.06 | 0.83 | 3.96 |
| 2 | 0.991 | 0.475 | 4ja3 | 0.87 | 2.46 | 0.86 | 2.76 | 0.88 | 2.22 | 0.88 | 2.22 | 0.88 | 2.38 |
| | | | 4ja4 | 0.83 | 3.40 | 0.81 | 3.84 | 0.84 | 3.04 | 0.84 | 3.06 | 0.84 | 3.12 |
| | | | 6n3i | 0.95 | 1.43 | 0.96 | 1.14 | 0.94 | 1.75 | 0.94 | 1.57 | 0.94 | 1.61 |
| 3 | 0.920 | 1.938 | 4ja3 | 0.84 | 3.00 | 0.84 | 2.99 | 0.76 | 4.28 | 0.85 | 2.86 | 0.85 | 2.87 |
| | | | 4ja4 | 0.80 | 4.05 | 0.80 | 4.12 | 0.79 | 3.93 | 0.81 | 3.96 | 0.81 | 3.85 |
| | | | 6n3i | 0.97 | 0.89 | 0.97 | 0.85 | 0.80 | 4.64 | 0.97 | 0.91 | 0.97 | 0.94 |
| 4 | 0.990 | 0.573 | 4ja3 | 0.86 | 2.77 | 0.86 | 2.72 | 0.83 | 3.37 | 0.85 | 3.16 | 0.87 | 2.64 |
| | | | 4ja4 | 0.82 | 3.56 | 0.83 | 3.57 | 0.80 | 4.28 | 0.81 | 4.02 | 0.83 | 3.35 |
| | | | 6n3i | 0.97 | 1.14 | 0.97 | 1.12 | 0.96 | 1.15 | 0.97 | 0.96 | 0.97 | 1.21 |
| 5 | 0.989 | 0.593 | 4ja3 | 0.87 | 2.50 | 0.86 | 2.84 | 0.88 | 2.35 | 0.91 | 1.79 | 0.87 | 2.54 |
| | | | 4ja4 | 0.83 | 3.22 | 0.82 | 3.76 | 0.84 | 3.10 | 0.86 | 2.25 | 0.83 | 3.24 |
| | | | 6n3i | 0.97 | 1.18 | 0.98 | 0.96 | 0.96 | 1.31 | 0.94 | 1.80 | 0.97 | 1.16 |
| 6 | 0.990 | 0.277 | 4ja3 | 0.94 | 0.68 | 0.94 | 0.61 | 0.94 | 0.64 | 0.93 | 0.82 | 0.94 | 0.70 |
| | | | 4ja4 | 0.92 | 1.52 | 0.94 | 1.20 | 0.92 | 1.48 | 0.90 | 1.60 | 0.91 | 1.52 |
| | | | 6n3i | 0.86 | 3.30 | 0.84 | 3.80 | 0.86 | 3.29 | 0.87 | 3.00 | 0.86 | 3.21 |
| 7 | 0.994 | 0.279 | 4ja3 | 0.94 | 0.79 | 0.94 | 0.88 | 0.95 | 0.55 | 0.94 | 0.65 | 0.94 | 0.73 |
| | | | 4ja4 | 0.92 | 1.51 | 0.90 | 1.57 | 0.93 | 1.26 | 0.92 | 1.37 | 0.92 | 1.46 |
| | | | 6n3i | 0.88 | 3.59 | 0.88 | 3.92 | 0.86 | 3.55 | 0.86 | 3.62 | 0.87 | 3.73 |

Protein Accession Number/Region: Mutated Residues

| | |
|---|---|
| 1 | WP_001097274 |
| 2 | N/C split (220/276): 24, 28, 31, 32, 35, 36, 50, 54, 55, 57, 58, 59, 61, 62, 65, 66, 68, 69, 169, 172, 175, 176, 179, 180, 182, 183, 186, 187, 297, 298, 299, 301, 305, 311, 312, 315, 316, 318, 319, 323, 325, 326, 329, 416, 417, 419, 420, 423, 424, 427, 428, 431, 432, 433 |
| 3 | N/C split (220/276) C-term only: 297, 298, 299, 301, 305, 311, 312, 315, 316, 318, 319, 323, 325, 326, 329, 416, 417, 419, 420, 423, 424, 427, 428, 431, 432, 433 |
| 4 | N/C split (220/276) N-term only: 24, 28, 31, 32, 35, 36, 50, 54, 55, 57, 58, 59, 61, 62, 65, 66, 68, 69, 169, 172, 175, 176, 179, 180, 182, 183, 186, 187 |
| 5 | G58W, L315W |
| 6 | Region 154-173: 5, 10, 14, 18, 21, 22, 24, 25, 147, 148, 156, 158, 163, 166, 167, 169, 170, 171, 172, 326, 329 |
| 7 | Region 327-341: 172, 328, 329, 332, 336, 337, 338, 339, 340, 381, 384, 385, 390, 465, 467 |

The TM and RMSD were calculated using residues 9-465.
The average TM and RMSD are the average values for the five models against each other in a single run (10 total comparisons).
Numbers in blue are the best TM or RMSD score for that structure.
Numbers in red are the best TM or RMSD score for that structure, but not for that model indicating that this structure may not be represented in the models.
The residues were mutated to Alanine unless otherwise noted.

| Supplemental Table 5: GLUT5 | | | | | | | | | | | | | |
|---|---|---|---|---|---|---|---|---|---|---|---|---|---|
| | Average | | | Model 1 | | Model 2 | | Model 3 | | Model 4 | | Model 5 | |
| Run | TM | RMSD | PDB | TM | RMSD | TM | RMSD | TM | RMSD | TM | RMSD | TM | RMSD |
| 1 | 0.929 | 1.847 | 4yb9 | 0.85 | 3.07 | 0.97 | 0.87 | 0.97 | 0.83 | 0.85 | 2.94 | 0.87 | 2.58 |
| | | | 4ybq | 0.93 | 0.82 | 0.81 | 3.60 | 0.82 | 3.44 | 0.94 | 0.82 | 0.93 | 1.01 |
| 2 | 0.988 | 0.628 | 4yb9 | 0.94 | 1.50 | 0.94 | 1.65 | 0.94 | 1.50 | 0.94 | 1.41 | 0.93 | 1.55 |
| | | | 4ybq | 0.80 | 3.76 | 0.80 | 3.77 | 0.80 | 3.76 | 0.81 | 3.56 | 0.84 | 2.97 |
| 3 | 0.993 | 0.466 | 4yb9 | 0.84 | 3.39 | 0.84 | 3.43 | 0.83 | 3.56 | 0.87 | 2.95 | 0.82 | 3.62 |
| | | | 4ybq | 0.93 | 0.98 | 0.93 | 1.04 | 0.93 | 0.98 | 0.93 | 1.28 | 0.94 | 0.87 |

| Protein Accession Number/Region: Mutated Residues | |
|---|---|
| 1 | NP_001315548 |
| 2 | From Run 1-1; N/C split (210/275):<br>32, 76, 78, 79, 82, 83, 86, 87, 89, 90, 93, 94, 98, 139, 140, 143, 144, 147, 148, 151, 152, 156, 157, 158, 159, 160, 161, 163, 164, 167, 168, 171, 172, 175, 178, 182, 183, 184, 315, 316, 318, 319, 322, 323, 325, 326, 329, 330, 333, 336, 337, 341, 392, 393, 397, 400, 401, 405, 408, 409, 412, 413, 416, 417, 419, 420, 423, 424, 428, 465, 466, 467, 468, 470 |
| 3 | From Run 1-2; N/C split (210/275):<br>32, 36, 40, 41, 47, 65, 68, 69, 71, 72, 75, 76, 77, 79, 80, 82, 83, 86, 140, 143, 164, 167, 171, 172, 175, 178, 181, 182, 183, 184, 187, 297, 298, 299, 300, 301, 302, 311, 315, 316, 318, 319, 322, 323, 325, 326, 329, 333, 392, 417, 419, 420, 423, 424, 427, 428, 431, 432, 435 |

The TM and RMSD were calculated using residues 19-462.
The average TM and RMSD are the average values for the five models against each other in a single run (10 total comparisons).
Numbers in blue are the best TM or RMSD score for that structure.
The residues were mutated to Alanine.

| Supplemental Table 6: Mhp1 | | | | | | | | | | | | | |
|---|---|---|---|---|---|---|---|---|---|---|---|---|---|
| | Average | | | Model 1 | | Model 2 | | Model 3 | | Model 4 | | Model 5 | |
| Run | TM | RMSD | PDB | TM | RMSD | TM | RMSD | TM | RMSD | TM | RMSD | TM | RMSD |
| 1 | 0.963 | 0.842 | 2jln | 0.99 | 0.77 | 0.98 | 0.88 | 0.98 | 0.95 | 0.91 | 2.20 | 0.94 | 1.67 |
| | | | 2x79 | 0.86 | 2.19 | 0.87 | 2.21 | 0.86 | 2.31 | 0.94 | 1.20 | 0.91 | 1.83 |
| | | | 4d1b | 0.98 | 0.64 | 0.98 | 0.64 | 0.97 | 0.68 | 0.93 | 1.85 | 0.96 | 1.25 |
| 2 | 0.995 | 0.370 | 2jln | 0.92 | 1.90 | 0.91 | 1.98 | 0.92 | 1.91 | 0.92 | 1.98 | 0.92 | 1.81 |
| | | | 2x79 | 0.92 | 1.62 | 0.92 | 1.68 | 0.91 | 1.86 | 0.92 | 1.52 | 0.90 | 1.98 |
| | | | 4d1b | 0.94 | 1.47 | 0.94 | 1.62 | 0.94 | 1.50 | 0.94 | 1.65 | 0.94 | 1.36 |
| 3 | 0.996 | 0.307 | 2jln | 0.91 | 1.99 | 0.91 | 2.02 | 0.91 | 1.99 | 0.91 | 1.98 | 0.91 | 2.12 |
| | | | 2x79 | 0.92 | 1.47 | 0.92 | 1.69 | 0.92 | 1.67 | 0.92 | 1.64 | 0.93 | 1.36 |
| | | | 4d1b | 0.93 | 1.61 | 0.93 | 1.66 | 0.93 | 1.62 | 0.93 | 1.62 | 0.93 | 1.78 |
| 4 | 0.983 | 0.654 | 2jln | 0.97 | 0.92 | 0.96 | 1.07 | 0.94 | 1.38 | 0.94 | 1.44 | 0.93 | 1.65 |
| | | | 2x79 | 0.86 | 2.50 | 0.85 | 2.58 | 0.88 | 2.12 | 0.88 | 2.08 | 0.90 | 1.71 |
| | | | 4d1b | 0.97 | 0.81 | 0.96 | 0.78 | 0.95 | 1.05 | 0.95 | 1.16 | 0.94 | 1.45 |

| Protein Accession Number/Region: Mutated Residues | |
|---|---|
| 1 | D6R8X8 |
| 2 | Region 159-170 (TM5): 26, 31, 34, 35, 38, 161, 164, 165, 167, 168, 169, 230, 233, 311, 312, 316, 320 |
| 3 | Region 159-170 (TM5) Complement: 26, 31, 34, 35, 38, 230, 233, 311, 312, 316, 320 |
| 4 | Region 352-368 (TM 9/10): 109, 113, 116, 117, 119, 123, 146, 353, 359, 363, 365, 366, 367, 423 |

The TM and RMSD were calculated using residues 11-466.

The average TM and RMSD are the average values for the five models against each other in a single run (10 total comparisons).

Numbers in blue are the best TM or RMSD score for that structure.

Numbers in red are the best TM or RMSD score for that structure, but not for that model indicating that this structure may not be represented in the models.

The residues were mutated to Alanine unless otherwise noted.

| Supplemental Table 7: LAT1 | | | | | | | | | | | | | |
|---|---|---|---|---|---|---|---|---|---|---|---|---|---|
| | Average | | | Model 1 | | Model 2 | | Model 3 | | Model 4 | | Model 5 | |
| Run | TM | RMSD | PDB | TM | RMSD | TM | RMSD | TM | RMSD | TM | RMSD | TM | RMSD |
| 1 | 0.997 | 0.285 | 7dsq | 0.87 | 2.98 | 0.87 | 2.90 | 0.87 | 2.88 | 0.87 | 2.95 | 0.88 | 2.78 |
| | | | 6irs | 0.96 | 1.09 | 0.96 | 1.13 | 0.96 | 1.11 | 0.96 | 1.15 | 0.95 | 1.28 |
| 2 | 0.995 | 0.356 | 7dsq | 0.92 | 2.12 | 0.92 | 2.06 | 0.91 | 2.37 | 0.90 | 2.57 | 0.90 | 2.42 |
| | | | 6irs | 0.93 | 1.52 | 0.93 | 1.56 | 0.94 | 1.31 | 0.94 | 1.27 | 0.94 | 1.44 |
| 3 | 0.963 | 0.752 | 7dsq | 0.97 | 1.10 | 0.97 | 1.09 | 0.90 | 1.69 | 0.94 | 1.48 | 0.96 | 1.44 |
| | | | 6irs | 0.86 | 2.81 | 0.86 | 2.87 | 0.86 | 2.72 | 0.90 | 2.19 | 0.89 | 2.22 |
| 4 | 0.969 | 0.887 | 7dsq | 0.96 | 1.31 | 0.89 | 2.03 | 0.91 | 2.26 | 0.93 | 1.79 | 0.89 | 2.68 |
| | | | 6irs | 0.87 | 2.65 | 0.95 | 1.49 | 0.94 | 1.53 | 0.92 | 1.80 | 0.95 | 1.25 |

| Protein Accession Number/Region: Mutated Residues | |
|---|---|
| 1 | NP_003477 |
| 2 | Region 188-206 (TM5): 64, 190, 193, 194, 197, 200, 203, 204, 205, 330, 333, 334, 337, 338, 340, 341, 344, 372 |
| 3 | Region 382-406 (TM9/10): 70, 73, 74, 95, 135, 138, 139, 140, 142, 143, 145, 146, 147, 150, 168, 172, 176, 180, 245, 248, 251, 252, 383, 384, 385, 387, 388, 390, 391, 393, 394, 395, 396, 397, 398, 399, 400, 401, 402, 403, 404, 405, 439, 447, 450, 451, 457, 458, 461, 462, 464, 465 |
| 4 | Region 52-63 (TM1a): 55, 56, 58, 59, 60, 259, 260, 277, 281 |

The TM and RMSD were calculated using residues 51-507.

The average TM and RMSD are the average values for the five models against each other in a single run (10 total comparisons).

Numbers in blue are the best TM or RMSD score for that structure.

The residues were mutated to Alanine.

| Supplemental Table 8: MCT1 | | | | | | | | | | | | | |
|---|---|---|---|---|---|---|---|---|---|---|---|---|---|
| | Average | | | Model 1 | | Model 2 | | Model 3 | | Model 4 | | Model 5 | |
| Run | TM | RMSD | PDB | TM | RMSD | TM | RMSD | TM | RMSD | TM | RMSD | TM | RMSD |
| 1 | 0.954 | 0.277 | 7da5 | 0.91 | 1.09 | 0.91 | 1.06 | 0.91 | 1.03 | 0.91 | 1.07 | 0.91 | 1.03 |
| | | | 7ckr | 0.74 | 4.22 | 0.74 | 4.16 | 0.74 | 4.24 | 0.75 | 4.04 | 0.74 | 4.22 |
| 2 | 0.953 | 0.399 | 7da5 | 0.76 | 3.74 | 0.78 | 3.45 | 0.76 | 3.64 | 0.76 | 3.72 | 0.76 | 3.71 |
| | | | 7ckr | 0.91 | 0.93 | 0.91 | 1.07 | 0.91 | 1.02 | 0.91 | 0.89 | 0.91 | 1.00 |
| 3 | 0.948 | 0.576 | 7da5 | 0.70 | 4.91 | 0.71 | 4.55 | 0.73 | 4.24 | 0.72 | 4.37 | 0.70 | 4.90 |
| | | | 7ckr | 0.89 | 1.49 | 0.91 | 1.22 | 0.92 | 0.92 | 0.91 | 0.97 | 0.88 | 1.69 |
| 4 | 0.867 | 2.161 | 7da5 | 0.74 | 4.23 | 0.75 | 4.00 | 0.69 | 4.83 | 0.76 | 3.72 | 0.74 | 4.30 |
| | | | 7ckr | 0.90 | 1.19 | 0.92 | 0.88 | 0.71 | 4.36 | 0.91 | 0.88 | 0.90 | 1.21 |

| Protein Accession Number/Region: Mutated Residues | |
|---|---|
| 1 | NP_001159968 |
| 2 | N/C split (194/261): 38, 41, 42, 44, 45, 46, 54, 55, 56, 59, 60, 63, 64, 66, 67, 70, 71, 151, 155, 156, 158, 159, 162, 163, 166, 281, 282, 283, 285, 286, 289, 295, 296, 299, 302, 303, 306, 310, 313, 395, 399, 402, 403, 404, 406, 407, 411, 413, 414, 419 |
| 3 | N/C split (194/261) C-term only: 38, 41, 42, 44, 45, 46, 54, 55, 56, 59, 60, 63, 64, 66, 67, 70, 71, 151, 155, 156, 158, 159, 162, 163, 166 |
| 4 | N/C split (194/261) N-term only: 281, 282, 283, 285, 286, 289, 295, 296, 299, 302, 303, 306, 310, 313, 376, 379, 395, 399, 402, 403, 404, 406, 407, 411, 413, 414, 419 |

The TM and RMSD were calculated using residues 16-449.
The average TM and RMSD are the average values for the five models against each other in a single run (10 total comparisons).
Numbers in blue are the best TM or RMSD score for that structure.
The residues were mutated to Alanine.

| Supplemental Table 9: MurJ | | | | | | | | | | | | | |
|---|---|---|---|---|---|---|---|---|---|---|---|---|---|
| | Average | | | Model 1 | | Model 2 | | Model 3 | | Model 4 | | Model 5 | |
| Run | TM | RMSD | PDB | TM | RMSD | TM | RMSD | TM | RMSD | TM | RMSD | TM | RMSD |
| 1 | 0.992 | 0.589 | 5t77 | 0.97 | 0.75 | 0.96 | 1.11 | 0.97 | 0.59 | 0.96 | 1.15 | 0.96 | 0.90 |
| | | | 6nc9 | 0.71 | 5.88 | 0.75 | 5.23 | 0.73 | 5.66 | 0.74 | 5.43 | 0.72 | 5.64 |
| 2 | 0.891 | 1.770 | 5t77 | 0.70 | 5.05 | 0.83 | 3.4 | 0.69 | 5.41 | 0.69 | 5.31 | 0.78 | 4.75 |
| | | | 6nc9 | 0.86 | 1.71 | 0.82 | 3.73 | 0.89 | 1.15 | 0.88 | 1.34 | 0.87 | 2.61 |
| 3 | 0.949 | 0.681 | 5t77 | 0.90 | 2.38 | 0.89 | 2.75 | 0.90 | 2.52 | 0.82 | 3.57 | 0.84 | 2.62 |
| | | | 6nc9 | 0.78 | 4.46 | 0.80 | 4.24 | 0.78 | 4.52 | 0.86 | 3.21 | 0.75 | 4.22 |
| 4 | 0.941 | 0.639 | 5t77 | 0.75 | 5.05 | 0.79 | 4.14 | 0.76 | 4.94 | 0.71 | 5.05 | 0.72 | 5.22 |
| | | | 6nc9 | 0.92 | 1.46 | 0.90 | 2.41 | 0.93 | 1.46 | 0.88 | 1.19 | 0.86 | 1.47 |
| 5 | 0.964 | 0.598 | 5t77 | 0.73 | 4.56 | 0.69 | 5.84 | 0.69 | 5.77 | 0.69 | 5.60 | 0.69 | 5.79 |
| | | | 6nc9 | 0.87 | 1.81 | 0.87 | 1.06 | 0.88 | 1.14 | 0.87 | 1.30 | 0.89 | 1.13 |

| Protein Accession Number/Region: Mutated Residues | |
|---|---|
| 1 | WP_012580438 |
| 2 | N/C split (243): 22, 25, 26, 28, 29, 30, 31, 32, 33, 34, 38, 39, 42, 230, 231, 234, 235, 238, 239, 241, 242, 248, 249, 251, 252, 255, 316, 317, 319, 320, 321, 322, 326, 371, 374, 375, 378, 382, 383, 384, 385, 386, 387, 390 |
| 3 | Region 223-234: 50, 53, 54, 57, 59, 108, 109, 225, 226, 229, 230, 231, 233, 371, 374, 375 |
| 4 | Region 248-255: 25, 28, 29, 33, 34, 38, 42, 249, 250, 251, 252, 253, 317, 322, 326, 329, 330, 333, 334, 337, 387 |
| 5 | Region 230-241: 42, 43, 46, 107, 108, 109, 110, 111, 112, 113, 231, 233, 234, 235, 236, 237, 238, 239, 240, 246, 247, 371, 374, 375, 378, 382, 383, 387, 390 |

The TM and RMSD were calculated using residues 4-470.
The average TM and RMSD are the average values for the five models against each other in a single run (10 total comparisons).
Numbers in blue are the best TM or RMSD score for that structure.
The residues were mutated to Alanine.

| Supplemental Table 10: PfMATE | | | | | | | | | | | | | |
|---|---|---|---|---|---|---|---|---|---|---|---|---|---|
| | Average | | | Model 1 | | Model 2 | | Model 3 | | Model 4 | | Model 5 | |
| Run | TM | RMSD | PDB | TM | RMSD | TM | RMSD | TM | RMSD | TM | RMSD | TM | RMSD |
| 1 | 0.997 | 0.241 | 3vvn | 0.98 | 0.53 | 0.99 | 0.56 | 0.99 | 0.57 | 0.98 | 0.59 | 0.98 | 0.64 |
| | | | 6fhz | 0.74 | 5.23 | 0.75 | 5.12 | 0.74 | 5.22 | 0.74 | 5.48 | 0.73 | 5.56 |
| 2 | 0.996 | 0.297 | 3vvn | 0.78 | 4.25 | 0.79 | 4.36 | 0.75 | 4.26 | 0.79 | 4.21 | 0.79 | 4.28 |
| | | | 6fhz | 0.93 | 1.12 | 0.94 | 1.08 | 0.93 | 1.13 | 0.93 | 1.03 | 0.94 | 1.03 |
| 3 | 0.992 | 0.378 | 3vvn | 0.79 | 4.41 | 0.78 | 4.40 | 0.78 | 4.45 | 0.79 | 4.34 | 0.77 | 4.43 |
| | | | 6fhz | 0.93 | 1.12 | 0.93 | 1.31 | 0.95 | 1.13 | 0.93 | 1.30 | 0.92 | 1.58 |
| 4 | 0.982 | 0.656 | 3vvn | 0.80 | 4.10 | 0.80 | 4.03 | 0.80 | 4.03 | 0.75 | 4.67 | 0.80 | 4.08 |
| | | | 6fhz | 0.93 | 1.19 | 0.93 | 1.50 | 0.92 | 1.39 | 0.90 | 1.95 | 0.92 | 1.49 |

| Protein Accession Number/Region: Mutated Residues | |
|---|---|
| 1 | WP_011011833 |
| 2 | N/C split (272): 12, 13, 14, 16, 17, 19, 20, 21, 23, 24, 27, 28, 31, 32, 35, 67, 68, 69, 71, 72, 73, 75, 76, 78, 79, 80, 82, 83, 84, 86, 87, 95, 96, 99, 100, 103, 161, 237, 238, 239, 241, 242, 245, 246, 249, 250, 253, 288, 291, 292, 294, 295, 296, 298, 299, 300, 302, 303, 305, 306, 308, 309, 310, 315, 318, 319, 322, 330, 387, 388, 390 |
| 3 | N/C split (272) N-term only: 12, 13, 14, 16, 17, 19, 20, 21, 23, 24, 27, 28, 31, 32, 35, 67, 68, 69, 71, 72, 73, 75, 76, 78, 79, 80, 82, 83, 84, 86, 87, 95, 96, 99, 100, 103, 161 |
| 4 | N/C split (272) C-term only: 237, 238, 239, 241, 242, 245, 246, 249, 250, 253, 288, 291, 292, 294, 295, 296, 298, 299, 300, 302, 303, 305, 306, 308, 309, 310, 315, 318, 319, 322, 330, 387, 388, 390 |

The TM and RMSD were calculated using residues 17-454.

The average TM and RMSD are the average values for the five models against each other in a single run (10 total comparisons).

Numbers in blue are the best TM or RMSD score for that structure.

The residues were mutated to Alanine.
